# Alpha globin variation in the long-tailed macaque suggests malaria selection

**DOI:** 10.1101/2020.10.21.344853

**Authors:** C.L. Faust, F. Rangkuti, S. G. Preston, A. Boyd, P. Flammer, B. Bia, N. J. Rose, F. B. Piel, A. L. Smith, A.P. Dobson, S. Gupta, B. S. Penman

**Author notes:** CORRESPONDANCE TO (BSP), (SG) and (ALS).

## Abstract

Human haemoglobin variants, such as sickle, confer protection against death from malaria; consequently, frequencies of such variants are often greatly elevated in humans from malaria endemic regions. Among non-human primates, the long-tailed macaque, *Macaca fascicularis*, also displays substantial haemoglobin variation. Almost all *M. fascicularis* haemoglobin variation is in the alpha globin chain, encoded by two linked genes: *HBA1* and *HBA2*. We demonstrate that alpha globin variation in *M. fascicularis* correlates with the strength of malaria selection. We identify a range of missense mutations in *M. fascicularis* alpha globin and demonstrate that some of these exhibit a striking *HBA1* or *HBA2* specificity, a pattern consistent with computational simulations of selection on genes exhibiting copy number variation. We propose that *M. fascicularis* accumulated amino acid substitutions in its alpha globin genes under malaria selection, in a process that closely mirrors, but does not entirely converge with, human malaria adaptation.

## Introduction

It is well established that certain human haemoglobin mutations reach high frequencies due to the protection they offer against death from malaria (Haldane 1949; Allison 1954a; Allison 1954b; Taylor, et al. 2012). Non-human primate species also host malaria parasites (Coatney, et al. 1971), but the question of whether non-human primate haemoglobins are under malaria selection is unresolved. In typical adult humans, over 97% of haemoglobin is made up of beta globin transcribed from the *HBB* gene, and alpha globin transcribed from two genes in the alpha globin cluster (*HBA1* and *HBA2*) that encode identical proteins (Weatherall and Clegg 2001b). Haemoglobin in other primates appears to be broadly similar, although gene duplications and deletions of *HBA* occur frequently as a result of unequal crossing over (Hoffman, et al. 2008). The common ancestor of old world primates, a group that includes macaques, is likely to have had three *HBA* genes in its alpha globin cluster (Hoffman, et al. 2008). 587 amino acid changes have been reported in human *HBB*; 89 in human *HBA1* and 150 in *HBA2* (Patrinos, et al. 2004). Only three of these amino acid substitutions reach significant frequencies in human populations: haemoglobin S (that leads to sickle cell anaemia when inherited in the homozygous state), haemoglobin C and haemoglobin E (Weatherall and Clegg 2001a), and all are caused by mutations in *HBB*. The former two offer significant malaria protection (Taylor, et al. 2012); clinical studies to prove the same for haemoglobin E are lacking. In addition to amino acid substitutions, thalassaemic mutations affecting the rate of alpha or beta globin subunit production exist in humans. There is strong evidence that alpha thalassaemia protects against severe malaria (Taylor, et al. 2012); beta thalassaemia likely also offers protection, but far fewer studies have been carried out to confirm this.

Long-tailed macaques (*Macaca fascicularis*) display the highest level of haemoglobin variation reported in a non-human primate species (supplementary table 1). Extensive surveys of haemolysates during the 1960s-80s revealed that *M. fascicularis* populations possess a variety of adult haemoglobins that can be distinguished using starch-gel electrophoresis (Barnicot, et al. 1966) (fig. 1, supplementary table 2). Band “A” haemoglobin migrates similarly to human adult haemoglobin and is likely the wild-type haemoglobin as it is present in all populations of *M. fascicularis* and sister species of long-tailed macaques. Band “Q” haemoglobin migrates anodally to **A** at an alkaline pH. Changes in the alpha globin subunit are responsible for the difference between **A** and **Q** (Barnicot, et al. 1966), specifically **Q** differs from **A** by the substitution of aspartic acid for glycine at alpha globin position 71 (p. Gly71Asp) (Takenaka, et al. 1988). A third electrophoretic variant, “P”, has only been found in mainland Southeast Asian populations of *M. fascicularis*. **P** results from an as-yet-undefined amino acid change at position 15 or 16 of alpha globin (Barnicot, et al. 1966) and **P** has been observed to polymerize *in vitro*. Finally, a “minor” haemoglobin (“X”) that migrates more slowly than **A** at an alkaline pH has been observed in some studies of *M. fascicularis* (Barnicot, et al. 1966; Ishimoto, et al. 1970). **X** is distinguished from **A** by four variant sites, all in alpha globin (p. Asn9Lys, p. Trp14Leu, p. Gly19Arg and p. Gly71Arg) (Wade, et al. 1970). In contrast to the widespread variation in alpha globin, only one beta globin variant has been found in a single isolated population of *M. fascicularis* (Kawamoto, et al. 1984).

**Fig. 1.**
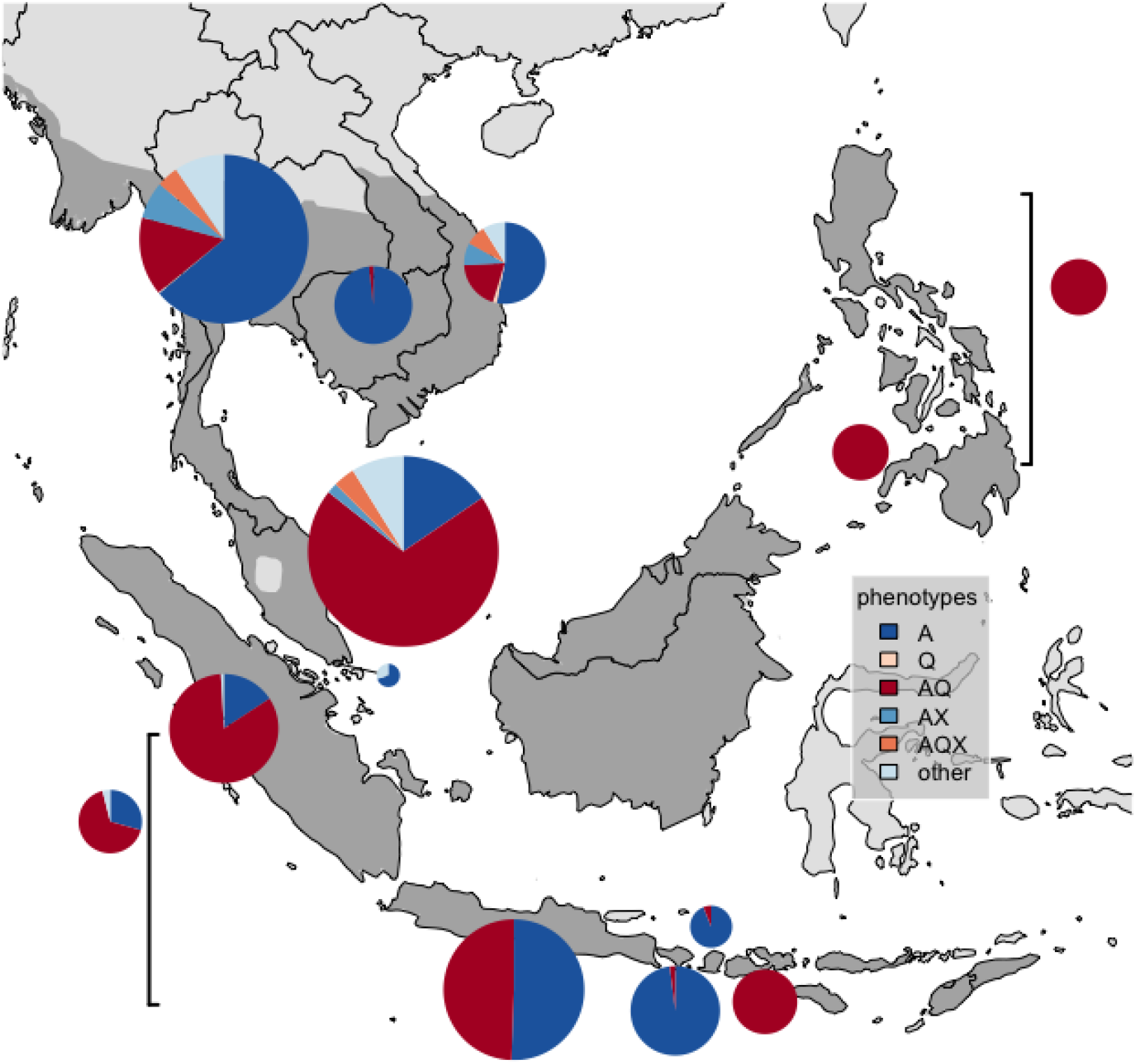
Map of haemoglobin variant phenotype frequencies. Surveys of haemoglobin were collected from published studies of long-tailed macaques, *Macaca fascicularis* (see supplementary table 2 for full references). The range of *M. fascicularis* is in dark grey (International Union for the Conservation of Nature 2019). Square brackets indicate surveys that come from Indonesia or the Philippines without an island specified (for the Philippines the only island that was specified in any survey was Mindanao). Radii of pie charts are determined by the sample size of macaques studied at that location (range: n = 10 for Singapore; n = 677 for Malaysia).

The first evidence for malaria selection acting on humans was the geographical association JBS Haldane observed between the presence of malaria disease and the presence of inherited blood disorders (Haldane 1949). Here, we demonstrate a geographical association between macaque haemoglobin variation and the presence of virulent macaque malarias, which provides a compelling rationale to extend Haldane’s malaria hypothesis to non-human primates. We further show, using parallel amplicon sequencing and population genetic simulations, that the *M. fascicularis* alpha globin cluster exhibits patterns which are consistent with selection.

## Results

### *M. fascicularis* alpha globin variation is more likely to be observed in the presence of virulent macaque malarias

For six locations where electrophoretic alpha globin phenotypes have been surveyed (fig. 1, supplementary table 2), macaque malaria surveys have also been carried out (Faust and Dobson 2015). These are Cambodia, Singapore, Thailand, Peninsular Malaysia and the Indonesian islands of Java and Bali (supplementary table 3, supplementary fig. 1). The **A** band occurs in every macaque population surveyed; thus we consider it to be ancestral. Unlike **P** and **Q, X** was not always examined in every study. This leads us to define **A** in the absence of **P** or **Q** to be a *non-variant alpha globin phenotype*, and any phenotype containing **Q**, **P** or both to be a *variant alpha globin phenotype*. Bayesian inference allows us to calculate the probability of observing at least one virulent macaque malaria species (*Plasmodium coatneyi* or *P. knowlesi*) in a region, and this estimate is used as a proxy for malaria selection (see Methods for virulent malaria justification). This Bayesian hierarchical model allows us to define the probability of observing a macaque with a variant alpha globin phenotype in regions with low or high malaria selection (Eq. 1, Methods, supplementary information section 1.2). In regions with low malaria selection, the probability of observing a long-tailed macaque with a variant alpha globin phenotype is 0.034 (95% credible interval (CI): 0.012, 0.053). In regions with high malaria selection, the probability of observing a long-tailed macaque with a variant alpha globin phenotype is much higher, 0.686 (95%CI: 0.660, 0.715). Sensitivity analyses demonstrate that lower probabilities of variant phenotypes are always found in areas that are unlikely to have virulent malarias, regardless of the specific proxy for malaria selection used (supplementary information section 1.2; supplementary figs. 2-4). There is, therefore, a geographical association between the presence of virulent malarias and variant alpha globin phenotypes in long-tailed macaque populations. The distribution of macaque haemoglobin variants is not simply explained by the heterozygosity of the macaque populations in these locations (supplementary information section 1.3, supplementary fig. 5); however we must acknowledge that it is not possible to reliably assess the heterozygosity of all the macaque populations in all the geographical locations of interest.

### Multiple unique HBA1 and HBA2 globin sequences are present in Indonesian long-tailed macaques

The two alpha globin genes (HBA1 and HBA2) in the*Macaca_fascicularis_*_5.0 genome (NC_022291.1) can be distinguished based on differences in their downstream sequences (for specific primers see Methods and SI). Out of a sample of 78 Indonesian *M. fascicularis* we successfully amplified and sequenced a 334 nucleotide region of both HBA1 and HBA2 from 77 animals. This 334 nucleotide region included exon 2 and parts of its flanking introns. We identified 13 unique HBA1 and 12 unique HBA2 globin sequences based on 24 variable sites (table 1). The majority of animals possessed 2 unique *HBA1* sequences (fig. 2A, supplementary fig. 6). For most animals we observed 3 unique *HBA2* sequences, but one macaque had 5 unique *HBA2* sequences (fig. 2A, supplementary fig. 7). These results are consistent with a single copy of *HBA1* and up to 3 copies of *HBA2* within the alpha globin gene cluster of these animals. From the proportion of reads each sequence contributed to the total (supplementary figs. 6-8), it would appear that some animals may possess more than 3 copies of *HBA1* or *HBA2* in their alpha globin clusters. We are reluctant to over-interpret the relative proportions of reads found, since it is possible that primers may have been biased towards amplifying certain sequences, but overall it seems likely that gene duplication of at least *HBA2*, and likely both *HBA1* and *HBA2* occurs within this population.

**Fig. 2.**
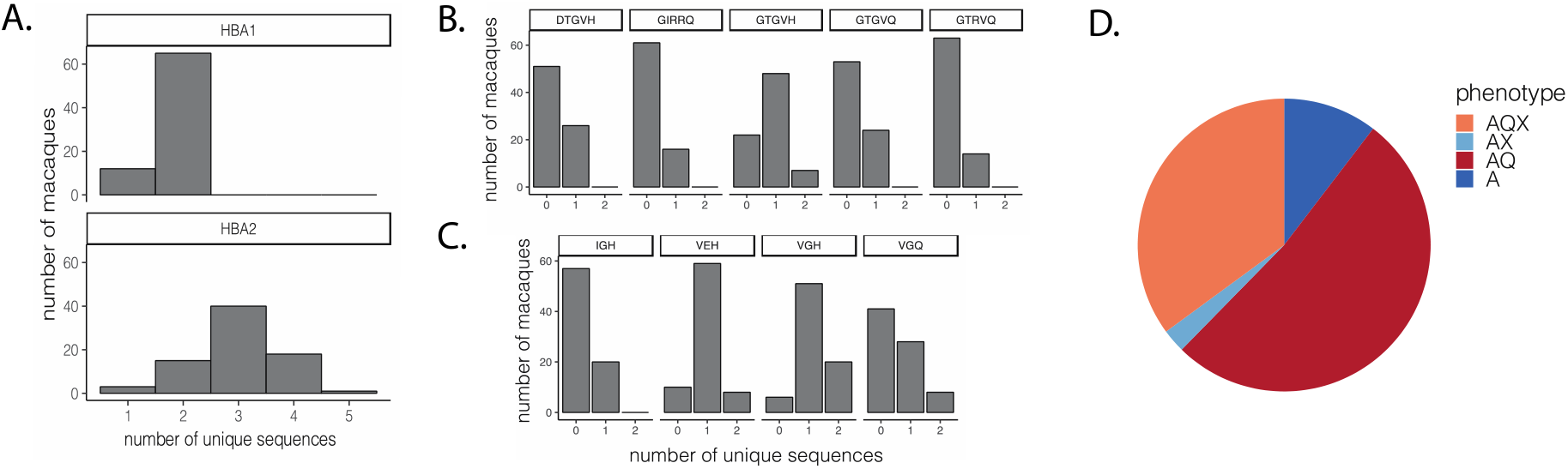
Distribution of *HBA1* and *HBA2* sequences observed in Indonesian *M. fascicularis*. A) The majority of long-tailed macaques possessed two unique *HBA1* sequences and three unique *HBA2* sequences. B) Five unique combinations of amino acid acids (variable sites were found at positions 57, 67, 71, 73, and 78) were observed amongst *HBA1* in the 77 long-tailed macaques. C) Four unique amino acid combinations (variable amino acid sites are listed at 55, 71 and 78) were found in *HBA2*. D) Predicted population frequency of electrophoretic phenotypes on the basis that **Q** can be generated by Gly71Glu and **X** by Gly71Arg.

**Table 1.**
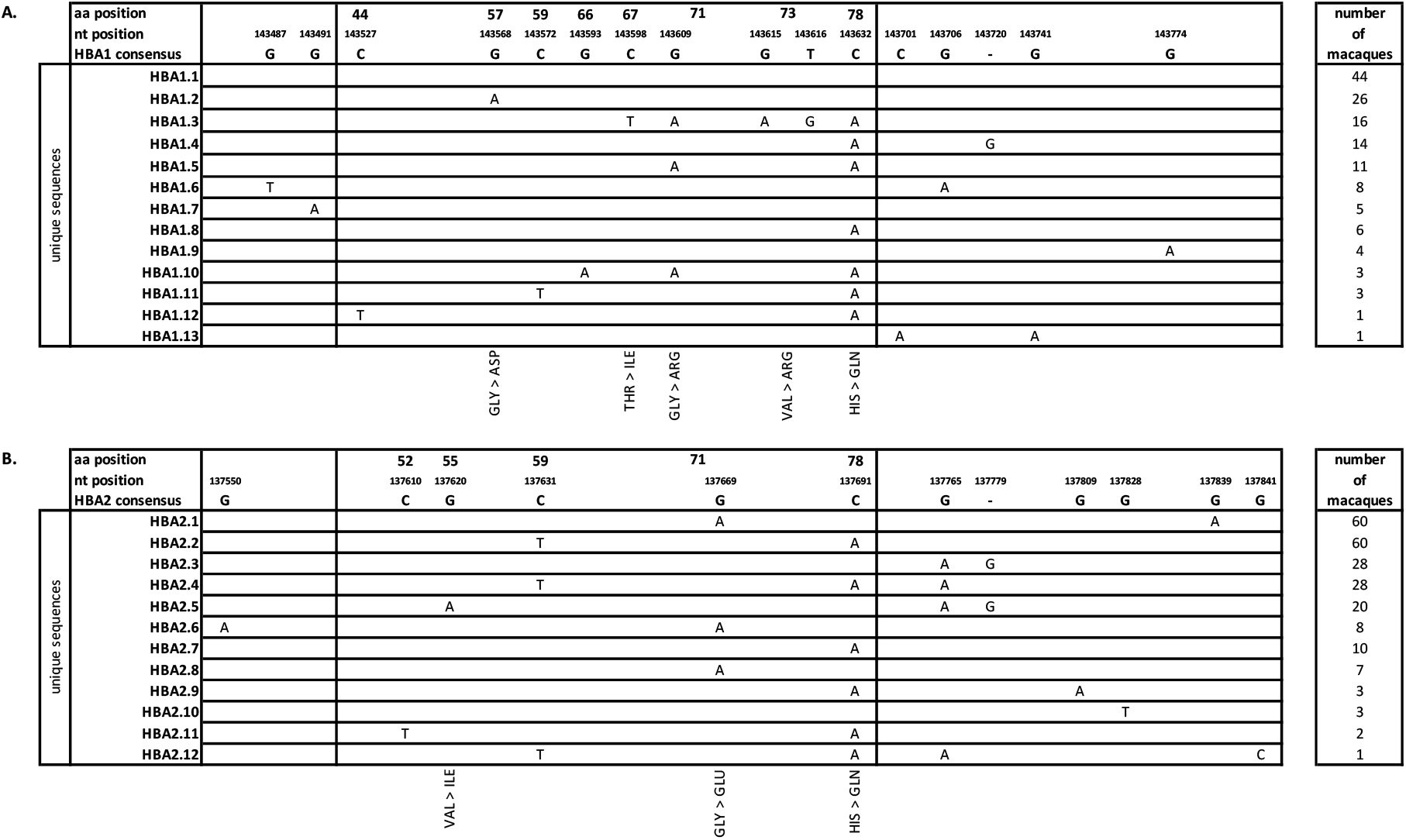
Unique *HBA1* and *HBA2* sequences in our population of Indonesian long-tailed macaques. The header row indicates the nucleotide and amino acid positions of each variant site in *M. fascicularis HBA1* (A) and *HBA2* (B). Nonsynonymous substitutions are indicated at the bottom of the table. The reference alpha globin sequences were taken from Chromosome 20 of a whole genome shotgun sequencing of *M. fascicularis*: Macaca_fascicularis_5.0 (NC_022291.1; Indonesian origin). The nucleotide position (nt pos) is the location along the reference sequence. Unique sequences are named for whether they were observed in *HBA1* or *HBA2*, and are also assigned a numerical identifier. The “number of macaques” column gives the frequency of animals in which the sequence is found. A 6 nucleotide sequence (ACGGGA) is present at positions 137834-137839 of the NC_022291.1 sequence (in the post exon 2 intron of HBA2) which we did not find in any of our samples. This appears to be a deletion in the intron of all our samples relative to the reference genome. Since this is not an example of variation within our samples we have not displayed it in table 1B.

Given our uncertainty over the exact number of copies of each of *HBA1* and *HBA2* present in each animal, it was not possible to predict the exact haplotypic combinations of *HBA1* and *HBA2* sequences present in each alpha globin cluster. However, some patterns are apparent, e.g. two unique *HBA2* globin sequences: HBA2.1 and HBA2.2 are always found together, and at similar proportions for each macaque (supplementary fig. 7) suggesting they may be linked on the same haplotype. In supplementary tables 6 and 7 we propose a potential set of haplotypes which could account for most of the genotypes in our sample. The most frequent haplotypes under this scheme are HBA1.1-HBA2.1-HBA2.2; HBA1.2-HBA2.3; HBA1.1-HBA2.5 and HBA1.3-HBA2.1-HBA2.2-HBA2.4.

### Amino acid position 71 displays specificity of SNPs between HBA1 and HBA2

We recorded seven nonsynonymous mutations across both *HBA1* and *HBA2* from this population of long-tailed macaques (table 1), of which Gly57Asp; Val73Arg; Gly71Arg and Gly71Glu had not previously been reported (fig. 2B,C). We observed two different substitutions at amino acid position 71, which has been shown to be the key amino acid site differentiating allozymes in electrophoretic studies (Takenaka, et al. 1988). All position 71 SNPs in *HBA1* sequences caused a change from glycine to arginine (Gly71Arg), and all position 71 SNPs in *HBA2* sequences caused a change from glycine to glutamic acid (Gly71Glu). Previous work identified a Gly71Asp substitution in **Q** bands in haemolysate from a single southern Sumatran *M. fascicularis* (Takenaka, et al. 1988). Both the previously observed amino acid change and the different *HBA2* substitution observed in our study involve a large negatively charged amino acid (glutamic acid or aspartic acid) replacing a small non-charged amino acid (glycine). We propose that these changes will give rise to similar phenotypic consequences, and that the Gly71Glu of *HBA2* is extremely likely to generate the **Q** band of *M. fascicularis* haemolysate (fig. 2D). The Gly71Arg change we observe in *HBA1* is one of the changes identified as characteristic of the **X** band (a so-called “minor haemoglobin” identified in some *M. fascicularis* – see Introduction). Other changes that are found in the **X** band occur in exon 1 (Wade, et al. 1970), and are beyond the scope of this analysis. If all the sequences we have identified are expressed, we predict the distribution of electrophoretic types within our sample to be as follows: **A**:8, **AQ**: 40, **AQX**:27, **AX**:2 (fig. 2D). Such a distribution of electrophoretic phenotypes is not unprecedented: populations with a high frequency of the **AQ** phenotype are known to exist in Indonesian populations of *M. fascicularis* (see supplementary table 2). **X** can be observed alongside the **A** and **Q** bands in *M. fascicularis* in other parts of its range (Barnicot, et al. 1966; Ishimoto, et al. 1970). Although **X** has not been reported in electrophoretic surveys of *M. fascicularis* from Indonesia, it is likely previous studies did not use protocols capable of detecting this variant (Kawamoto and Ischak 1981; Kawamoto, et al. 1984; Perwitasari-Farajallah, et al. 1999).

### Multiple peptide sequences are possible in HBA1 and HBA2

Of the nonsynonymous substitutions we found at amino acid positions other than 71, His78Gln and Thr67Ile have been previously reported to occur within *M. fascicularis* **A** band (Takenaka, et al. 1988). Thr67Ile is only found in *HBA1* sequences in our sample; His78Gln is found in both *HBA1* and *HBA2*. We identified two additional substitutions unique to *HBA1:* Gly57Asp and Val73Arg. Gly57Asp is likely to alter the electrophoretic properties of haemoglobin, by analogy with the same change in human alpha globin where it gives rise to the fast migrating Hb J-Norfolk (Baglioni 1962). Likewise, Val73Arg may give rise to similar electrophoretic properties as Gly71Arg change. Interestingly all 16 individuals displaying Val73Arg also had Gly71Arg, but how Val73Arg may relate to the X band of *M. fascicularis* haemoglobin is unclear. We also observed a novel Val55Ile substitution in *HBA2* in 20 individuals. The biochemical similarity of valine and isoleucine means it is possible that this change would not significantly alter the properties of haemoglobin.

Despite the range of substitutions observed among our samples, it was not possible to phylogenetically determine evolutionary relationships between these different *M. fascicularis* alpha globin exon 2 sequences (supplementary fig. 9). This is likely due to the fact that we had only a relatively short sequence of 334 nucleotides. The length of our sequence and the fact that only 10/68 (15%) of codons in exon 2 are variable means dn/ds ratios are unlikely to be able to provide reliable insights into whether positive selection is evident among these sequences (Anisimova, et al. 2002).

### Natural selection can maintain high frequency HBA1 or HBA2 specific polymorphisms

The HBA1 and HBA2 specificity of certain amino acid substitutions in our sample is a surprising finding given HBA paralogues typically have highly similar coding regions in primates (almost certainly a consequence of gene conversion)(Hoffman, et al. 2008). To understand whether natural selection can drive the *HBA1* or *HBA2* specificity of amino acid substitutions in *M. fascicularis*, we simulated *HBA1* and *HBA2* in a finite diploid population using an individual based model (see Methods). Mutations generating two different amino acid substitutions at the same site in alpha globin, one neutral and one potentially under selection, were able to enter the population via migration. Gene conversion (*c*) and reciprocal crossing over (*r*) could occur between *HBA1* and *HBA2* and the probabilities of each were varied (fig. 3A-C). Our model thus assumed the simulated population to be connected to a wider global population of *M. fascicularis*, acting as the source of haemoglobin variation.

**Fig. 3.**
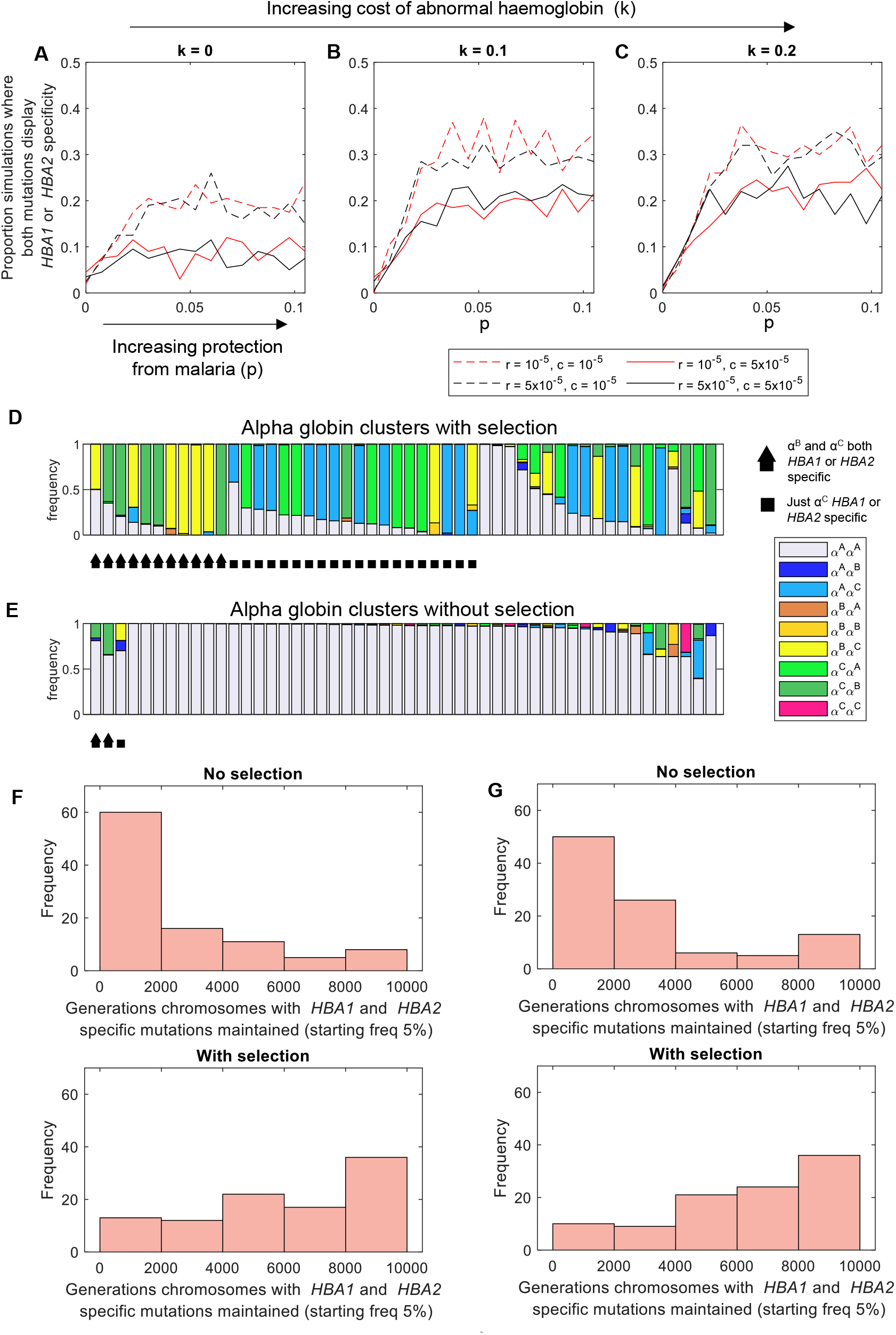
Selection increases the probability of observing *HBA1* or *HBA2* specific amino acid substitutions. Three alpha globin types are possible in the model: the ancestral type (α^A^); a neutral variant (α^B^) and a potentially selected variant (α^C^). Panels A-C illustrate how the properties of α^C^ affect the probability of observing a population genetic outcome in which both non ancestral types are present in the population, each associated with only *HBA1* or only *HBA2* (see Methods for a more detailed description of the thresholds used to define this state). *p* (x axis of each panel) is the protection against an environmental hazard such as malaria provided by having ≥1 copies of α^C^ in a genotype, and *k* (title of each panel) is the disadvantage (i.e. blood disorder cost) associated with having ≥3 copies of α^C^ in a genotype. The populations simulated for panels A-C all started with 100% α^A^ at both *HBA1* and *HBA2*. Two different gene conversion (c) and reciprocal crossing over (r) probabilities were used, as indicated in the legend. Each of these are probability of gene conversion or reciprocal crossing over affecting the gametes of an individual macaque (see Methods for justification). Other parameters were: N=10000, t=10000, m= 0.2 and d=0.05. 200 repeats were carried out at each combination of parameters. Panels (D and E) each visualise the distribution of chromosome types present at the end of 50 individual simulations. Each stacked bar represents one simulation and the relative proportions of the bands within each stacked bar indicate different possible chromosomes (see legend). If selection is included, k=0.2 and p = 0.03. If selection is not included k=0 and p=0. Other parameters in panels D and E were: r= 10^-5^, c= 10^-5^, N=10000, t=10000, m= 0.2 and d=0.05. Panels (F and G) display histograms of the number of generations for which at least 2 out of the 3 chromosomes α^C^α^A^, α^A^α^B^ and α^C^α^B^ were maintained at frequencies >2.5% in simulations of a closed population (m=0) where the starting frequencies of each chromosome were: 5% α^A^α^B^; 5% α^C^α^A^; 5% α^C^α^B^ and 85% α^A^α^A^. In panel F, r= 10^-5^ and c= 10^-5^ and in panel G r= 0 and c= 0. In both F and G, if selection is included, k=0.2 and p = 0.03; if selection is not included k=0 and p=0. Other parameters in F and G were N=10000, d=0.05, t=10000. 100 repeats were carried out at each combination of parameters.

The simultaneous maintenance of both mutations as *HBA1* or *HBA2* specific polymorphisms in the population is most likely when there is (i) an advantage to the state where some, but not all, alpha globin genes in a genotype encode the selected variant, and (ii) a cost to the state where more than two of the alpha globin genes in a genotype encode the selected variant (fig. 3A-C). *HBA1* or *HBA2* specificity of both mutations simultaneously is often (but not exclusively) achieved when both the neutral and the selected mutations are present on the same chromosome, and that chromosome is elevated to a high frequency (fig. 3D). Selection is also likely to elevate the selected variant alone in an *HBA1* or *HBA2* specific manner (fig. 3D). Without selection, we see fewer scenarios in which any mutations are present at high frequencies. Among those where mutations do reach higher frequencies there is no bias towards *HBA1* or *HBA2* specificity (fig. 3E).

Figure 3A-E assume a high level of gene flow with a large global *M. fascicularis* population continually providing new genetic diversity. In the absence of such a process, populations will, eventually, become fixed for a single alpha globin cluster (bearing the selected variant if selection is present). Figure 3F and 3G illustrate how long multiple alpha globin clusters bearing *HBA1* and *HBA2* specific mutations persist in the absence of gene flow (each mutant chromosome starting at a frequency of 5%). Selection extends the average time that multiple mutant chromosomes can coexist, because chromosomes bearing the selected mutation (some of which may also carry a neutral mutation) are preferentially maintained in the population.

We cannot be certain of the historical population size of Indonesian *M. fascicularis*, and whether or not the mutations we have found in our sample arose *de novo* in Indonesia or were imported from elsewhere. It is therefore not possible to calculate exactly how likely it is that the pattern in our sample arose under an entirely neutral model – but we can say that under two possibilities: frequent challenge with diverse mutations (fig. 3A-E) or an entirely closed system (fig. 3F-G), selection acting on at least some of the mutations makes the maintenance of multiple HBA1 or HBA2 specific mutations far more likely.

## Discussion

We have conducted the first DNA analysis of the alpha globin cluster of *M. fascicularis*. The most striking feature of our results is that several amino acid changes are limited to *HBA1* or *HBA2*, with a stark separation of two possible substitutions at alpha globin amino acid position 71. Gly71Arg and Gly71Glu both occur as part of more than one unique *HBA1* (Gly71Arg) or *HBA2* (Gly71Glu) sequence. This indicates that their *HBA1* and *HBA2* specificity is widespread in Indonesian *M. fascicularis*, and not a result of inbreeding in the colony. Our population genetic simulations show that such *HBA1* or *HBA2* specificity is more likely to be maintained over longer periods of time if at least one of these amino acid substitutions is under selection. Amino acid substitutions at alpha globin position 71 are associated with the electrophoretic phenotypes **A**, **AQ** and **AQX** explored in historical studies. Our geographical analysis showed that there is an association between variant haemoglobin electrophoretic phenotypes in *M. fascicularis* and the presence of virulent macaque malarias. We contend, therefore, that the most likely selective pressure to account for the *HBA1* and *HBA2* specificity of alpha globin variants in *M. fascicularis* is malaria selection.

The closest evolutionary relative of *M. fascicularis* is the rhesus macaque, *Macaca mulatta*. Data from a recently published whole genome sequencing study of *M. mulatta* (Xue, et al. 2016) allows us to analyse whether the polymorphisms we identified in *M. fascicularis* occur in its sister species. We were able to obtain partial alpha globin exon 2 sequences for 98 *M. mulatta* (supplementary information section 1.5). The only non-synonymous change detected in the *M. mulatta* samples was the His78Gln substitution also found in *M. fascicularis*, where it occurs in both *HBA1* and *HBA2* sequences (supplementary table 8). Unfortunately, the short reads used for whole genome sequencing make it impossible to distinguish *HBA1* from *HBA2* sequences in the *M. mulatta* samples. The reasons for the maintenance of His78Gln as a (possible) trans-species polymorphism are unclear. However, the fact that we observed none of the *HBA1* or *HBA2* specific amino acid changes belonging to *M. fascicularis* in the *M. mulatta* data shows that *HBA1* or *HBA2* specific amino acid substitutions are not necessarily a feature of macaque alpha globin generally. It has been noted that *M. mulatta* and *M. fascicularis* differ in their susceptibility to malaria, and that malaria itself may have driven their speciation (Wheatley 1980). Higher admixture of *M. fascicularis* with *M.mulatta* has been suggested to increase susceptibility to *P. cynomolgi* in breeding colonies (Zhang, et al. 2017). Our observations of *M. mulatta* and *M. fascicularis* alpha globin are consistent with these hypotheses.

Five other macaque species: *Macaca nemestrina*, *Macaca arctoides*, *Macaca assamensis, Macaca radiata*, and *Macaca sinica* possess variant haemoglobins which may be similar to *M. fascicularis* **Q** haemoglobin (fig. 4, supplementary table 9). *Macaca fascicularis* and *M. nemestrina* were the only macaques naturally found infected with the virulent parasites *P. knowlesi* and *P. coatneyi* (Eyles, et al. 1962) (supplementary table 10). However, recent surveys of *M. arctoides* from Thailand present evidence that these species are also naturally infected with *P. knowlesi* and *P. coatneyi* (Fungfuang, et al. 2020). *M. radiata*, the Bonnet macaque, is native to southwest India and is infected with three malaria species, including *P. fragile* (Ramakrishnan and Mohan 1962; Dissanaike, et al. 1965). *Plasmodium fragile* undergoes deep vascular schizogony like *P. coatneyi* and *P. knowlesi* and causes 33% mortality in intact *M. mulatta* (Eyles 1963; Coatney, et al. 1971). A range of different amino acid substitutions may be responsible for these different variant haemoglobins (see fig. 4 legend), but it is striking that a correlation between the presence of virulent malaria and variant macaque haemoglobins may extend beyond *M. fascicularis*.

**Fig. 4.**
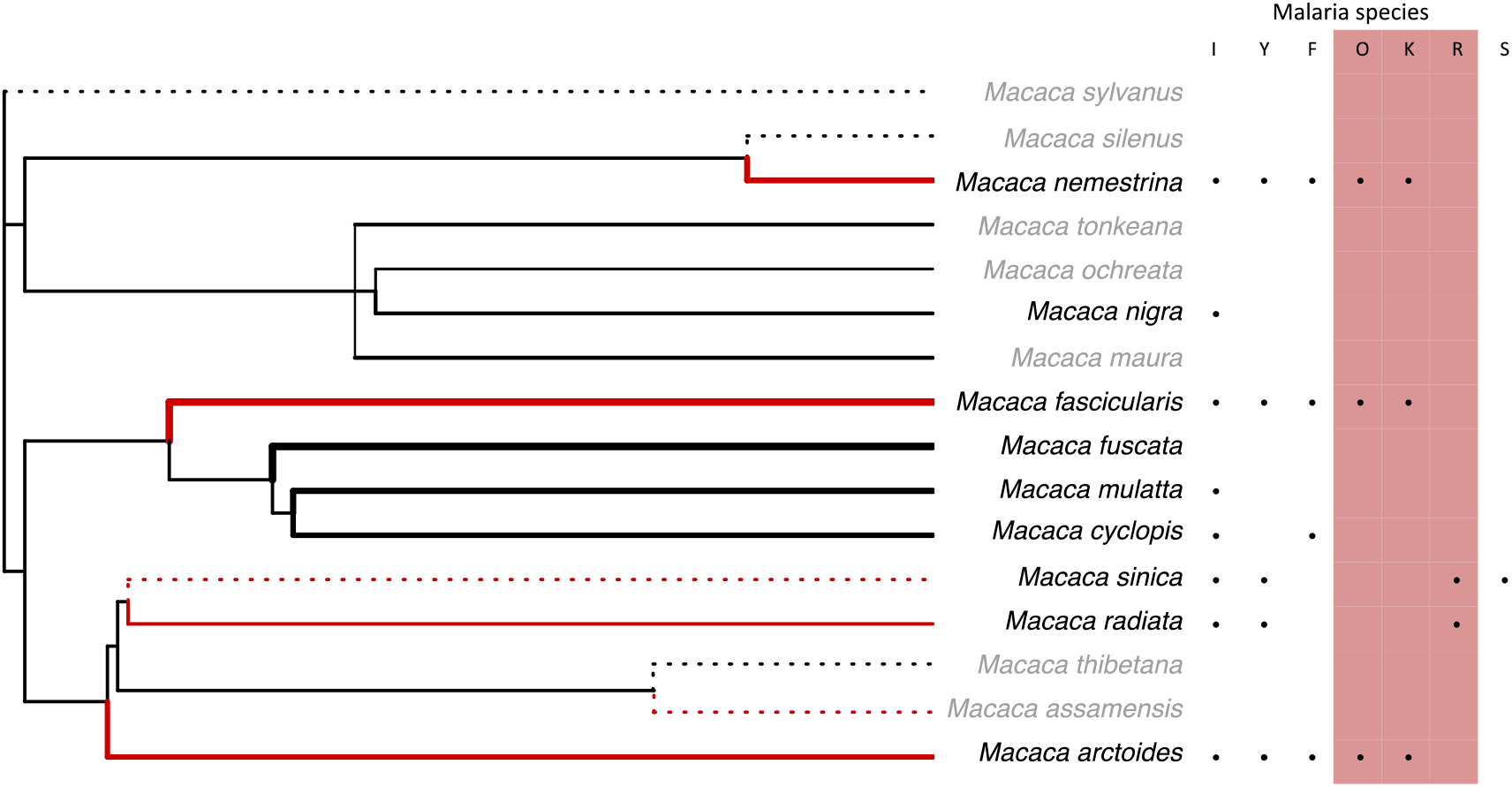
Major variant alpha globin phenotypes in macaques. The phylogeny is a species tree of macaques truncated from the mammalian super tree (Bininda-Emonds, et al. 2007). The branches of the species tree are annotated with information on the phenotypic variation in alpha globin from electrophoretic studies (supplementary table 9). The line type of the branches indicates whether we have (solid) or have not (dotted) been able to identify a population-level survey of alpha globin phenotypes in the literature (see supplementary table 9 for more details). The width of solid branches reflects the log of the total sample sizes of haemoglobin surveys (supplementary table 9)-thickest branch (*M. fuscata*) representing 2539 individuals and thinnest branch (*M. ochreata*) representing 17 individuals. Macaque species with variant alpha globin phenotypes are denoted by red branches and most (not *M. radiata*) are sympatric over part of their ranges. All variant alpha globin phenotypes for which we were able to identify the original source papers were reported to migrate similarly to **AQ** in *M. fascicularis* (original descriptions of *M. sinica* and *M. assamensis* haemoglobin variants could not be found – see supplementary table 9 for more details). The biochemical origins of haemoglobin variants in non-*fascicularis* species are as follows: alpha globin Gly71Asp substitutions have been found in *Macaca nemestrina* (Mahoney and Nute 1979; Takenaka, et al. 1988); an alpha globin Asp15Gly substitution is responsible for an **A/Q**-like polymorphism in *Macaca arctoides* (Oliver and Kitchen 1968; Maita, et al. 1985), and the same polymorphism is also found in *M. assamensis*. *M. sinica* possesses an Ala12Asp polymorphism. An unknown mutation results in a polymorphism electrophoretically similar to the **A/Q** polymorphism in *M. radiata* (Weiss, et al. 1973). The adjacent table of malaria species shows the parasites that have been found in each macaque species: *Plasmodium inui* (I), *P. cynomolgi* (Y), *P. fieldi* (F), *P. coatneyi* (O), *P. knowlesi* (K), *P. fragile* (R), and *P. simovale* (S) (supplementary table 10). The parasite species highlighted in red represent known virulent malarias, as defined in the Methods and Discussion. Macaque species names shown in grey are species that have not been sampled for malaria.

A protective effect of haemoglobin electrophoretic variant phenotypes could help explain puzzling results from experimental infection trials. *P. knowlesi* infection has a consistently mild course in *M. fascicularis* exported from the Philippines, whilst ‘Malayan’ animals directly exported from Singapore suffered fatal infection (Schmidt, et al. 1977). The **AQ** phenotype appears universal among *M. fascicularis* in the Philippines (supplementary table 2). Long-tailed macaques from Singapore, by contrast, are more likely to display an **A** band alone (proportion A band alone = 0.7, n =10) (Barnicot, et al. 1966) (fig. 1, supplementary table 2). A further study showed that *P. coatneyi* was fatal to splenectomized *M. fascicularis* from Mauritius, but not to splenectomized Philippine *M. fascicularis* (Migot-Nabias, *et al*. 1999). Mauritian animals are less likely to display variant alpha globin phenotypes (proportion variants = 0.09; n = 201) (Kondo, et al. 1993) than Philippine animals (proportion variants = 1.0; n = 118) (Ishimoto, et al. 1970; Ishimoto 1972), further suggesting that variant alpha globin phenotypes predict survival when infected with virulent malarias.

The infection study of *M. fascicularis* exported from the Philippines and Singapore also suggests a potential mechanism of protection by alpha globin variants. Only ring stage parasites were observed in blood of the lethally infected Singaporean animals, suggesting parasites were sequestering outside of peripheral circulation. However, all asexual development stages of the parasite were observed in the blood of the non-lethally-infected Philippine animals, suggesting parasite sequestration was less successful (Schmidt, et al. 1977). In humans, there is an association between sequestered parasite biomass and severe disease (Dondorp, et al. 2005). The parasite antigen PfEMP1 (*Plasmodium falciparum* erythrocyte membrane protein 1) is an important regulator of cytoadherence in the human parasite *Plasmodium falciparum*. *P. knowlesi* possesses a similar antigen called SICA (Schizont-infected cell agglutination antigen) (Brown and Brown 1965; Korir and Galinski 2006) although its role in cytoadherence is not well characterized. Human haemoglobinopathies can affect the expression of PfEMP1 (Fairhurst, et al. 2005; Cholera, et al. 2008), reducing cytoadherence. It is possible that macaque haemoglobin variants can affect the expression of *P. knowlesi* cytoadherence molecules, and that macaque red blood cells containing more than one major adult haemoglobin (as we expect to occur in all Philippine animals) are associated with reduced sequestration and improved health outcomes.

The extensive variation of *M. fascicularis* alpha globin is contrasted by just one beta globin amino acid substitution reported in *M. fascicularis* from Bali (Kawamoto, et al. 1984). *Macaca fascicularis* may therefore present an inversion of the situation in *Homo sapiens*, whose major malaria protective amino acid substitutions occur in beta globin, not alpha. Primate beta globin is encoded by a single gene in the beta globin cluster, whilst primate alpha globin is encoded by two or more linked genes in the alpha globin cluster. An alpha globin cluster can therefore contain two genes encoding structurally different alpha globin proteins. If such an alpha globin cluster becomes fixed in a population, the entire population might be able to express two or more different types of haemoglobin. It may be that this state is particularly advantageous against malaria. Since beta globin is typically encoded by just one gene in vertebrates, the equivalent situation is extremely unlikely to emerge for beta globin.

No human population has fully fixed a malaria protective haemoglobinopathy mutation, although the Tharu population of the Terai region of Nepal (a holoendemic malaria region) has come close, with alpha thalassaemia frequencies > 80%. The alpha thalassaemic mutation in question is a deletion of one of the two alpha globin genes in the alpha globin cluster. Deleting just one alpha globin gene in the cluster allows the remaining alpha globin gene to continue to support the function of the cell and the entire population can, theoretically, enjoy the same malaria protective phenotype if everyone is homozygous for the deletion. It may be, although the mutations are very different, that high frequencies of alpha thalassaemic deletions in the Tharu population and the fixation of the **AQ** phenotype among Philippine *Macaca fascicularis*, represent similar states of population adaptation to malaria.

An additional observation from our sequencing was that many or all of our studied population possessed more than two copies of alpha globin per chromosome, specifically at least two copies of *HBA2* in addition to *HBA1*. The sheer diversity of patterns observed when proportions of sequences are considered (supplementary figs. 6 –8) is also suggestive of variation in the number of copies of alpha globin. Previous studies have shown that triplication or quadruplication of alpha globin reaches high frequencies in certain *M. fascicularis* populations (Takenaka, et al. 1991; Takenaka, et al. 1993), and have noted varying proportions of variant haemoglobins in different samples (Barnicot, et al. 1966). If we allow for the possibility that an alpha globin cluster containing three alpha globin genes generates an excess of alpha globin chains, and that this has some malaria protective effect, then it is possible that *M. fascicularis* alpha globin copy number variation arose through its malaria protective advantage, and this advantage was subsequently enhanced by the incorporation of amino acid changes into some of the duplicated alpha globin genes. The contrasting routes that humans and *M. fascicularis* appear to have taken to achieve malaria protection may follow from the higher frequency of alpha globin deletions in humans (alpha thalassaemia), as opposed to alpha globin duplications in *M. fascicularis*.

There are alternative explanations for the observed patterns of genetic variation and fixation in alpha and beta globin genes in long-tailed macaques. Mitochondrial and low-coverage whole genome sequences demonstrate there is significant population structure between mainland and insular populations of long-tailed macaques (Tosi and Coke 2007; Kanthaswamy, et al. 2013; Yao, et al. 2020). While we find a compelling correlation between alpha globin phenotypic variation and malaria selection, it is possible that genetic drift on insular populations may explain some fixation of the **AQ** or **A** states, although we have checked for evidence of bottlenecks with available genetic data (supplementary figure 5). Our ability to test for positive selection (i.e. dn/ds ratios) within alpha globin is currently limited by only having short read sequences from a single population (Kryazhimskiy and Plotkin 2008). Long-read sequencing of globin genes across the population structure of longtailed macaques would allow the application of dn/ds ratio tests for positive selection on *M. fascicularis* alpha globin.

Haldane’s malaria hypothesis was developed to explain the geographical association between human malaria and heritable blood disorders. The malaria hypothesis has been validated with clinical evidence for a malaria protective effect of haemoglobinopathies in human populations. We find a geographical association between *M. fascicularis* alpha globin variant phenotypes and malaria selection, measured as the presence of virulent malaria species. Furthermore, we find that the specificity of amino acid variants to particular copies of alpha globin may be a signature of natural selection. Further research is required to prove that this selection is from malaria. Long read sequencing would provide higher quality data in order to correctly phase and assign haplotypes within this gene complex. This, combined with SNP panels to control for population structure and test for signatures of positive selection would provide more confidence in these findings. A significant challenge is demonstrating a benefit of variant alpha globins at the individual level in *M. fascicularis*. This could be done with experimental infections, but such experimentation carries significant ethical concerns.

It is becoming clear that there are many parallels between malaria resistance mechanisms among different vertebrate species. There are higher rates of adaptation in mammalian proteins that interact with *Plasmodium* species versus matched controls, and domains of alpha spectrin have been identified as potential sites for primate evolution in response to malaria (Ebel, et al. 2017). Examples of human malaria resistance traits with parallels in other species include sickle haemoglobin (a convergent form of sickle haemoglobin exists in deer (Esin, et al. 2017)) and FY variation (Duffy negativity confers human resistance to *Plasmodium vivax;* yellow baboon FY variation affects their susceptibility to malaria-like *Hepatocystis* parasites (Tung, et al. 2009)). As we add to the list of ways that different hosts have adapted the same proteins to combat the problem of malaria, we increase our potential to uncover biochemical similarities that advance our understanding of how each protective mechanism operates at the molecular level.

## Materials and Methods

### Geographical analyses

We identified twelve electrophoretic population surveys of *M. fascicularis* alpha globin from the literature (Barnicot, et al. 1966; Barnicot, et al. 1970; Ishimoto, et al. 1970; Ishimoto 1972; Nozawa, et al. 1977; Smith and Ferrell 1980; Kawamoto and Ischak 1981; Kawamoto, et al. 1984; Kawamoto, et al. 1989; Tomiuk 1989; Kondo, et al. 1993; Perwitasari-Farajallah, et al. 1999) (supplementary table 2). One study did not include sufficient geographical information so was excluded (Tomiuk 1989). Another survey was based on samples derived from an introduced population of Mauritian long-tailed macaques (Kondo, et al. 1993), where there is no active malaria transmission. We used the remaining surveys to conduct the geographic analyses. Specificity of sample origins ranged from troop-level latitude and longitude (Kawamoto, et al. 1984) to whole countries. Since our malaria selection likelihood calculation (detailed below) was at the regional level, we aggregated the alpha globin data by region (i.e. Sumatra Utara).

We sought to analyse a possible link between long-tailed macaque alpha globin phenotypic variation and malaria selection. This required us to develop a proxy for malaria selection across the range of long-tailed macaques. We used the likely presence of virulent malaria (*P. coatneyi* or *P. knowlesi*) in a region as our malaria selection proxy. We consider *P. coatneyi* and *P. knowlesi* to be the most virulent malarias infecting *M. fascicularis*, for the following reasons: (i) *P. coatneyi* and *P. knowlesi* have been shown to be capable of killing some, though not all, experimentally infected *M. fascicularis* (Schmidt, et al. 1977; Migot-Nabias, et al. 1999); (ii) unlike *P. cynomolgi, P. inui*, or *P. fieldi*, *P. coatneyi* and *P. knowlesi* cause lethal infections in the sister species of *M. fascicularis, M. mulatta* (Coatney, et al. 1971), and (iii) like *P. falciparum*, but unlike other macaque malarias, *P. knowlesi* and *P. coatneyi* undergo deep vascular schizogony – attachment of infected RBCs to the vascular endothelium–which may be associated with increased pathology (Desowitz, et al. 1969; Miller, et al. 1971). To assess the presence of *P. coatneyi* or *P. knowlesi* and malaria sampling effort across the range of *M. fascicularis*, we used a systematic review of publications that reported surveys of malaria in primates (Faust and Dobson 2015).

Using the number of long-tailed macaques surveyed for malaria and the number of macaques infected with virulent malarias (supplementary table 3), we fitted a probability density function (PDF; equation 1) for the presence of virulent malarias at each location using a beta distribution with a uniform prior:

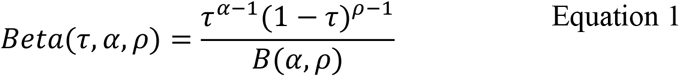

where α is the number of individuals with virulent malarias, ρ is the number of individuals without virulent malarias and 0 ≤ τ ≤ 1. The cumulative distribution function (CDF) for the probability density function (PDF) defined by equation 1 can then be used to determine a likelihood that virulent malaria is present or not present in a given locality by using a specific cutoff. We used a cutoff of 0.02 for the results reported in the main text, meaning we took a “likelihood of malaria being present” equal to the area under the PDF for a given region (equation 1) where the probability of observing a virulent malaria was >2% (and vice versa, the “likelihood of malaria not being present” was 1-the aforementioned area). A sensitivity analysis adjusting this cutoff is described in the supplementary methods (supplementary information section 1.2). We used a Metropolis-Hastings sampler with 100000 estimates, 20000 burn in and thinning every 100 to estimate the proportion of long-tailed macaques that had variant alpha globin phenotypes in areas with high malaria selection (high likelihood of virulent malarias) compared to low malaria selection (low likelihood of virulent malarias). The MCMC sampler was implemented in the *sampyl* package in Python 2.7 (Python Software Foundation 2010).

### Genetic characterization of HBA1 and HBA2 exon 2

Genomic DNA for 78 *Macaca fascicularis* from the UK National Institute for Biological Standards and Control (NIBSC) breeding colony was obtained from archived samples held by NIBSC. These samples were taken during historical health screening of the long-tailed macaques as part of standard colony management. The ancestors of these *M. fascicularis* were from Indonesia (further geographical specificity is unknown), and they have been bred in the UK for ~14 generations but still retain high MHC diversity (Mitchell, et al. 2012). *HBA1* and *HBA2* were amplified separately using gene-specific primers (supplementary table 5). For the initial PCR, reactions were run with 30ng template gDNA and initially heated to 98°C for 30sec, followed by 20 cycles of denaturation (98°C, 15 sec), annealing (70°C or 68°C, 20 sec), and extension (72°C, 30 sec), and then a final extension at 72°C for 2 minutes using Q5 High-Fidelity Polymerase (New England BioLabs). We conducted a nested PCR to sequence exon 2 (204bp; 68 amino acids) and flanking introns (5’ end: 9 bp; 3’ end: 121 bp) for both HBA amplicons, while adding a unique barcode to each sample: a 7 nucleotide sequence at the 5’ end of the primers formed a (7,4) Hamming barcode (Bystrykh 2012). 78 samples were multiplexed in a single MiSeq library in this way. The nested PCR was performed in a 50 μl solution containing polymerase master mix, 300 nM each of barcoded forward primer and barcoded reverse primer, and 1 μl of template DNA (supplementary table 2). Reactions were initially heated to 98 °C for 30 sec, followed by 10 cycles of 98 °C for 15 s, 68 °C for 20 s, 72 °C for 30 s. Reactions were completed at 72 °C for 2 min. Barcoded PCR products were pooled and purified using a QIAquick PCR Purification Kit (Qiagen, Hilden, Germany) according to manufacturer’s protocol. Illumina library preparation was performed with this pool using the NEBNext Ultra kit (NEB, Ipswich, MA) according to the manufacturer’s instructions, with size selection. Final concentration was measured by qPCR using the NEBNext Library Quant Kit for Illumina (NEB) according to the manufacturer’s instructions. Libraries were diluted from 26.2 nM (*HBA1*) and 33.4 nM (*HBA2*) to a concentration of 4 nM and pooled for sequencing on a MiSeq (Illumina, San Diego, CA).

Illumina output was demultiplexed using custom scripts in Python 2.7 (Python Software Foundation 2010). Output was denoised and dereplicated using dada2 v1.2.1 (Callahan, et al. 2016), and further manipulated using data.table v1.10.4 (Dowle and Srinivasan 2017) in R v3.2 (R Core Team 2015). Unique sequences that were found at a frequency less than 2% of an individual’s total reads were assumed to be PCR errors and were excluded from this analysis.

### Population genetic model

#### Model structure

We set up an individual based model of a diploid population of constant size *N*. The alpha globin cluster consists of two linked alpha globin genes (*HBA1* and *HBA2*) Each individual in the model thus posseses 4 globin genes, 2 in a cluster inherited from their mother and 2 in a cluster inherited from their father. The ancestral alpha globin type is designated type α^A^. Two further alpha globin types, α^B^ and type α^C^, represent possible (and mutually exclusive) amino acid changes in alpha globin. Every generation there is probability *m* that a single new migrant of a randomly generated genotype replaces an existing member of the population. The aforementioned randomly generated genotype consists of four globin types, randomly sampled with replacement from α^A^, α^B^ or α^C^, linked into two alpha globin clusters to create a diploid genotype. α^B^ is always a neutral variant, no more or less fit than α^A^. α^C^ may be under selection, generated by two processes. Firstly, *d* is the probability of any individual dying before reproducing in a given generation as a consequence of an environmental hazard (e.g. *d* could represent the burden of malaria on the population). Individuals with 1 or more genes encoding α^C^ in their genotype have a reduced probability of dying due to this hazard, such that their probability of dying from the hazard is equal to (1-*p*)*d*. Secondly, α^C^ may be associated with a blood disorder. Individuals with 3 or more genes encoding α^C^ in their genotype have probability *k* of dying before reproducing. Parameters *p* and *k* thus tune the advantages and disadvantages of genotypes containing globin type α^C^. If p=0 and k= 0 then α^C^ becomes a neutral variant. Every generation, those individuals which survive the environmental hazard and the potential blood disorder cost go on to form the parents of the next generation. During the reproduction step, randomly chosen pairs of parents each produce a single offspring genotype generated according to Mendelian inheritance until the required population size of *N* is reached. Reciprocal crossing over between *HBA1* and *HBA2* (i.e. the swapping of a maternal *HBA1* with a paternal *HBA1* between the two clusters so that each *HBA2* ends up linked to an alternate *HBA1*) takes place in each individual with probability *r*. Gene conversion, defined here as the conversion of one randomly chosen sequence within a genotype to match a randomly chosen sequence from the other three alpha globin sequences present in that genotype, takes place in each individual with probability *c*. The population evolves for *t* generations in each simulation.

In figure 3, a population genetic outcome with *HBA1* or *HBA2* specific mutations is defined as one in which:

- α^B^ and α^C^ are present in the population, each accounting for at least 5% of alpha globin sequences overall. This is to ensure that any *HBA1* or *HBA2* specificity was not a function of a mutation only being present at a very low frequency.
- At least 98% of α^B^ sequences in the population occur at *HBA1* or *HBA2* only, and likewise at least 98% of α^C^ sequences occur at *HBA1* or *HBA2* only.

Code used to implement this model is provided in ‘ AlphaGlobinPopulationGeneticModel.c’ at github.com/cfaustus/macaque_workspace.

#### Choice of parameters

We simulated a population size (*N*) of 10000. We simulated 10000 generations of evolution (t=10000). Longer and larger simulations were not possible due to computational limitations, so it was not practical to simulate the mutations arising entirely *de novo* within the population at a realistic rate. To get around this limitation we considered scenarios in which mutations arrive in the population at random, assumed to be generated in a wider global population of *M. fascicularis* (fig. 3A-E), and simulations in which genetic diversity is introduced at the beginning of the simulation, and the stability of that diversity considered over time, (fig. 3D-E).

We tested three possible values for the probability of gene conversion (*c*) and reciprocal crossing over (*r*) in our simulations: 0, 10^-5^ and 5×10^-5^. These were chosen based on the rate of unequal crossing in the human alpha globin cluster during meiosis (a study of human sperm observed 10^-5^ unequal crossing over events per sperm (Lam and Jeffreys 2007), thus a probability of 10^-5^ per meiosis event). Although our model is not simulating unequal crossing over, this is the best estimate we have for a reasonable alpha globin recombination rate.

## Supplementary Material

<u>Supplementary</u> Information includes supplementary text sections 1.1 – 1.5, supplementary tables 1-10, and supplementary figures 1-9.

The raw amplicon sequencing data of *Macaca fasciciularis* generated in this study have been deposited in the Sequence Read Archive (BioProject ID: PRJNA639946) under the accession numbers: SRR12404495-SRR12404572 (HBA1) and SRR12404678-SRR12404755 (HBA2)). Code associated with this research is available at github.com/cfaustus/macaque_workspace.

## Acknowledgements

This work was supported by a Princeton-Oxford Collaborative Grant; the Wellcome Trust (BSP grant 096063/Z/11/Z); the Truman Foundation (CLF); National Defense Science and Engineering (CLF); the Schlumberger Foundation (FR); the Oxford Centre for Islamic Studies (FR); the Royal Society (SG) and the European Research Council (SG, grant DIVERSITY). The funders had no role in study design, data collection and analysis, decision to publish, or preparation of the manuscript.

