## Supplementary Information for "Alpha globin variation in the long-tailed macaque suggests malaria selection"

#### 1. Supplementary Methods and Results

1.1 Alpha globin and malaria surveys in *Macaca fascicularis*
1.2 Sensitivity analyses of *M. fascicularis* alpha globin diversity and malaria prevalence
1.3 Heterozygosity within *M. fascicularis* populations surveyed
1.4 PCR amplification and sequencing
1.5 Bioinformatic analyses of *M. mulatta* alpha globin reads from the Sequence Read Archive.

#### 2. Supplementary Tables

Supplementary table 1. Haemoglobin variation across the Primate Order
Supplementary table 2. Alpha globin phenotypic survey data
Supplementary table 3. The prevalence of *Plasmodium* species infection among *M. fascicularis*
Supplementary table 4. Heterozygosity of *Macaca fascicularis* populations from different locations at non-globin loci
Supplementary table 5. Primers for amplifying *M. fascicularis HBA1* and *HBA2*
Supplementary table 6. Proposed *M. fascicularis* alpha globin haplotypes present in our sample
Supplementary table 7. Haplotype assignments and parental or sibling relationships within our sample
Supplementary table 8. Summary of *Macaca mulatta* exon 2 sequences for sites which are variable
Supplementary table 9. Haemoglobin surveys of the *Macaca* genus
Supplementary table 10. Occurrence of *Plasmodium* species in *Macaca*

#### 3. Supplementary Figures

Supplementary Figure 1. Malaria surveys in long-tailed macaques
Supplementary Figure 2. Robustness of the virulence analysis
Supplementary Figure 3. Varying haemoglobin surveys included in the analysis
Supplementary Figure 4. Non-A phenotype proportions in regions where minor haemoglobins were tested
Supplementary Figure 5. The relationship between the frequency of variant haemoglobins and the heterozygosity of different *M. fascicularis* populations
Supplementary Figure 6. Individual long-tailed macaque *HBA1* sequences
Supplementary Figure 7. Individual long-tailed macaque *HBA2* sequences
Supplementary Figure 8. The minimum proportion of any unique *HBA1* or *HBA2* sequence per animal
Supplementary Figure 9. Phylogenetic relationships between *M. fascicularis* alpha globin nucleotide sequences.

### 1. Supplementary Methods and Results

#### 1.1 Alpha globin and malaria surveys in *Macaca fascicularis*

##### 1.1.1 Interpreting historical alpha globin surveys

To make comparisons across all surveys, we required data at the alpha globin phenotype level. However, three of the studies listed in Supplementary table 2 (Kawamoto and Ischak 1981; Kawamoto, et al. 1984; Kawamoto, et al. 1989) reported allele frequencies rather than phenotypes, assuming a chromosomal arrangement based on the work of Smith and Ferrell (Smith and Ferrell 1980). Unfortunately this scheme did not address the so-called "minor haemoglobin" X noted in the main text. The results in the aforementioned studies were reported as allele frequencies where the alpha 1 locus was assumed to display alleles A or P ('1' or '4') and the alpha 2 locus was assumed to contain either a null allele ('0') or Q ('2'). To obtain phenotypic level data we back-transformed the data from these three studies. In a system assumed to contain only the A and the AQ chromosomes, where the true frequency of the AQ chromosome is *z*, the expected proportion of animals with the AQ phenotype (*p*) is given by equation S1.

$$p = 2(1 - z)z + z^2 \qquad \text{Equation S1}$$

We assume that the 'square root method' means z is related to p by equation S2.

$$z = \frac{2 \pm \sqrt{4 - 4p}}{2} \qquad \text{Equation S2}$$

We assumed 'z' is reported in each paper as the frequency of allele '2' at the alpha 2 locus. To obtain the AQ phenotype frequencies reported in Supplementary table 2 we applied Equation S1. Two troops of long-tailed macaques studied by Kawamoto, et al. (1984) and Kawamoto, et al. (1989) contained a low frequency of P haemoglobin (one troop in Sumatra Utara and one in Southern Thailand). Since the zymograms supplied in previous reports (Kawamoto and Ischak 1981; Kawamoto, et al. 1982) did not include any AQP or P individuals, we assumed that any 'P' haemoglobin must have occurred in the context of an 'AP' phenotype. We therefore assumed that the number of AP individuals in each survey must be equal to 2*N**y*, where y is the reported frequency of allele '4' at the alpha 1 locus and N is the number of animals in the sample. Only 7 animals displayed the AP phenotype in the two troops.

##### 1.1.2 Details of historical macaque malaria surveys

Findings for *M. fascicularis* across different geographical regions are summarized in supplementary table 3 and come from a systematic review of simian malaria (Faust and Dobson 2015). The presence of *P. coatneyi* or *P. knowlesi* in any given region required morphological or molecular identification. We did not consider molecular identification with SSU rRNA as sufficient evidence for the presence of a species, as the use of this gene can have low specificity when distinguishing *Plasmodium* species unless developmental stage is controlled and gene duplication is taken into account (Nishimoto, et al. 2008; Singh and Divis 2009).

#### 1.2 Sensitivity analyses of *M. fascicularis* alpha globin diversity and malaria prevalence

For the analyses described in the main text, we used a cutoff of 0.02 to determine the probability of virulent macaque malaria being present in a given locality (see main text Methods). In supplementary figure 2 we illustrate the effect of increasing this cutoff to 0.04. At the higher cutoff, there is still a substantial difference in the predicted frequency of variant alpha globin phenotypes between macaque populations exposed to virulent malarias and those not exposed to virulent malarias. For better comparison with the main text results, in all other sensitivity analyses presented here we use the cutoff of 0.02.

To obtain the predicted frequencies of variant alpha globin phenotypes reported in the main text, observations of long-tailed macaque alpha globin phenotypes from 6 countries and islands were used (Bali, Cambodia, Java, Singapore, Thailand, Peninsular Malaysia). These locations were chosen because they also had corresponding malaria surveys that reported *Plasmodium* presence to the species level. However, only including samples where malaria surveys had been conducted in the same country led us to exclude many of the haemoglobin surveys that were conducted. Therefore, we also conducted several additional analyses varying the geographic coverage of phenotype surveys to test the robustness of the initial patterns observed.

Malaria surveys in Mindanao and Sumatra did not report species identification for parasites found in long-tailed macaques. If we assume the Mindanao and Sumatra parasites are all non-virulent or all virulent, then we are able to include haemoglobin surveys from these locations in the analyses. Regardless of whether we assume Mindanao and Sumatra parasites to be all non-virulent or all virulent the distribution of posterior estimates does not change much compared to those reported in the main text (supplementary fig. 3A). This may be because the numbers of macaques surveyed for haemoglobin from the extra locations are relatively small (Mindanao, n= 59; Sumatra, n =222), compared to number of long-tailed macaques used to obtain the probabilities reported in the main text (n=1851). We also ran the analysis using all haemoglobin surveys of long-tailed macaques from areas where malaria is endemic, where, for regions where there are no macaque malaria surveys, we used data from the geographically closest malaria survey to give us a probability of observing virulent parasites. For Vietnam, Cambodia was used as a proxy. For macaque samples listed as only ‘Indonesian’, we used virulent malaria estimates from the entirety of Indonesia. This shifts the probability of observing variant phenotypes in the presence of virulent malarias slightly higher than that reported in the Main text (supplementary fig. 3B).

The main text and above scenarios consider the frequencies of haemoglobin variants identified by “major” electrophoresis bands, since those have been recorded for all the studies we identified. A subset of studies did record the presence of “minor” haemoglobin bands, including the X band, which, as noted in the main text, is caused by an amino acid change that is found only in the *HBA1* gene in our Indonesian samples – a potential hallmark of malaria selection. To examine the relationship between malaria selection and the probability of observing any variant haemoglobin, including haemoglobin X, we limited the dataset to surveys that looked for minor haemoglobins (these are surveys from Singapore, Thailand, Peninsular Malaysia and Vietnam). There are no malaria data surveys from Vietnam, so our final analysis was restricted to only five surveys conducted in three regions. We used the same likelihoods of virulent malaria as in the main text but now estimated the probability of observing any variant phenotype, including minor haemoglobin variants. The geographic distribution is heavily restricted (all adjoining locations on the Isthmus of Kra). The posterior estimates are shifted to higher probabilities compared to analyses only considering major haemoglobin phenotypes, but there is still a clear pattern that virulent malarias are associated with a higher probability of observing variant haemoglobins (supplementary fig. 4).

#### 1.3 Heterozygosity within *M. fascicularis* populations surveyed

We have shown that *M. fascicularis* populations where virulent malarias are present tend to possess higher frequencies of variant alpha globin electrophoresis phenotypes. However, *M. fascicularis* populations are known to exhibit substantial regional genetic structure (Tosi and Coke 2007; Kanthaswamy, et al. 2013; Yao, et al. 2020). Is it possible that the pattern we observe is not a consequence of malaria selection, but instead reflects greater *M. fascicularis* genetic diversity in regions which happen to also possess virulent malarias?

Supplementary figure 5 collates regional *M. fascicularis* heterozygosity data for a range of different loci, plotted against the proportion of each population displaying variant alpha globin (as defined in the main text). There is no clear pattern that a higher probability of observing variant haemoglobins corresponds to higher or lower heterozygosity in the population. In regions where virulent malarias are present, a wide range of heterozygosity can be observed at non globin loci. Of the two regions without any virulent malarias (Bali and Cambodia), which also have low frequencies of haemoglobin variants, Bali is a low heterozygosity insular population, but Cambodia is a high heterozygosity mainland population.

#### 1.4 PCR amplification and sequencing

For all PCR reactions, we used Q5® High-Fidelity DNA Polymerase in a 50 µl reaction with 30ng of template genomic DNA. To separate *HBA1* and *HBA2*, two separate PCR reactions were run to amplify *HBA1* and *HBA2*, using primers F3F-F8R and F3F-F4R to amplify each locus, respectively (supplementary table 5). The following PCR conditions were used to amplify DNA and minimize amplification of PCR error: an initial denaturation at 98°C for 30 seconds, followed by 20 cycles of denaturation (98°C, 15 sec), annealing (20 sec), and extension (72°C, 30 sec), and then a final extension at 72°C for 2 minutes. Annealing temperature for *HBA1* primers was 70°C, whereas *HBA2* primers were annealed at 68°C. After checking for amplification on 1% agarose gel stained with ethidium bromide, amplified DNA was subjected to a second PCR reaction with barcoded primers to amplify exon 2 and flanking introns (product size 374). This PCR reaction used MM2-S3R primers and had an initial denaturation at 98°C for 30 seconds, followed by 10 cycles of denaturation (98°C, 15 sec), annealing (68°C, 20 sec), and extension (72°C, 30 sec), and then a final extension at 72°C for 2 minutes.

*HBA1* and *HBA2* sequences present in individual long-tailed macaques are detailed in Table 1 of the main text and Supplementary figures 6 and 7. This diversity was observed even after excluding unique sequences where the read depth was less than 2% of the individuals' total reads. The average number of reads per individual for *HBA1* was 9140.71 and the average number of reads per individual for *HBA2* was 4205.47. One sample (MF_51) had only 29 reads for *HBA1* and 9 reads for *HBA2.* This sample was removed, and 77 individuals had a minimum of 300 reads per locus. If we removed the 2 individuals that had less than 1000 reads for *HBA1* and *HBA2* (MF1_42, Mf1_51) and an additional sample (Mf1_59) that also had less than 500 reads in *HBA2*, the distribution of predicted population phenotypes does not substantially change. Therefore, these samples with at least 300 reads were included in Supplementary figures 6 and 7 and in the analysis in the main text.

#### 1.5 Bioinformatic analyses of *M. mulatta* alpha globin reads from the Sequence Read Archive

##### 1.5.1 Methodology

A 2016 study carried out whole genome sequencing on 133 rhesus macaques (*M mulatta*) from multiple research facilities (Xue, et al. 2016). The data from this study are deposited in the sequence read archive (SRA) under accession PRJNA251548. We used this data to examine whether any of the alpha globin amino acid substitutions identified in our data are also present in *M. mulatta*.

The reads from each animal in the aforementioned 2016 study have already been mapped to a macaque whole genome shotgun sequence in which alpha globin appears only once. Exon 2 of alpha globin is located from positions 136747 to 136950 of the CM000306.1 chromosome 20 shotgun sequence (also known by the obsolete name NC_007877.1). We wrote a bespoke Python script to download the reads which mapped to this region, plus a surrounding +/- 100 bp (i.e. positions 136647 to 137050 of

CM000306.1) for each of the relevant runs in the Xue study from the SRA website in fastq format (see supplementary file MulattaReads.py). The supplementary material of the 2016 study lists biosample accession numbers for the animals in the study (SAMN02981228 etc), which we matched to run accession numbers (SRR1951019 etc) using information from the SRA website, in order to download animal-specific reads.

We used bowtie2(Langmead and Salzberg 2012) and samtools(Li and Durbin 2009) to produce an indexed alignment of *M. mulatta* reads for each animal. We aligned the reads to a reference sequence for *M. mulatta* alpha globin exon 2 and a surrounding +/- 100 bp region (Mmul_8.0.1 genome assembly, chromosome 20, positions 133708-134111). Finally, we used PySamStats (Miles 2018) to count the different nucleotides present at each site in exon 2 in the alignment for each animal. We filtered out low quality base calls by including the requirement "min_baseq = 20". Having established the numbers of high quality A, T, C or G calls at each site in alpha globin exon 2, we used the following rules to decide whether the site exhibited variation in a given animal. By allowing the inclusion of sites with a low read depth (4 and above) we maximised the information we were able to obtain from the somewhat patchy data.

If the read depth at a given site was <=20 reads, we applied the following thresholds:

- If there are <4 reads, designate the site "N", to indicate missing data.
- If there are >=3 reads indicating a single base, and <=1 read for any other base, identify the site as monomorphic for the highest frequency base.
- If there are >=2 reads for two different bases, and <=1 read for the other two bases, identify that that site as dimorphic for the two highest frequency bases, using standard ambiguity codes (A or T = W; A or G = R; A or C = M; T or G = K; T or C = Y; C or G = S).
- If there are >=2 reads for three or more different bases, give the site a "cannot identify" code Z

If the read depth at a given site was > 20 reads, we used the following cut offs:

- >85% reads indicating a single base, and <=5% for any of the other bases identifies the site as monomorphic for the highest frequency base.
- >5% of the reads for each of two bases, and the remaining bases represented by <=5% of reads, identifies that that site as dimorphic for the two highest frequency bases.
- >5% of reads for each of three or more different bases leads to us giving the site a "cannot identify" code Z (distinct from the missing data case "N").

The file MulattaReads.py included in the GitHub repository for this publication provides the code to perform these operations.

##### 1.5.2 Results

Using the method described in section 1.5.1, we obtained partial alpha globin exon 2 sequences for 98 out of the 133 macaques in Xue *et al*'s study. 32/98 animals had a mean read depth of >20 across exon 2; 71/98 had a mean read depth of >4. The remaining animals lacked coverage for part of exon 2, but had enough coverage in other regions of exon 2 to allow us to draw some conclusions based on the rules described in section 1.5.1.

We excluded any individual nucleotide substitutions which appeared in just one animal, on the basis that this variation could have been due to sample-specific PCR errors and/or our relatively lenient methods to identify variation. 12 potential substitutions were excluded in this way, distributed across 9 different macaques. The remaining sites that varied in *M. mulatta* were a synonymous substitution at amino acid position 69, GCC/GCT (Ala), which we did not observe in any of our *M. fascicularis* samples (main text), and a non-synonymous substitution at amino acid position 78, CAC/CAA (His>Gln), which we did observe in both *HBA1* and *HBA2 M. fascicularis* sequences (main text). Supplementary table 8 summarises the *M. mulatta* genotypes we observed at the aforementioned two variable *M. mulatta* sites. Supplementary table 8 also shows the number of *M. mulatta* samples for which we observed invariant genotypes at sites we found to be variable in *M. fascicularis*.

*HBA1* and *HBA2* can only be distinguished by a region downstream of exon 3, which is >500 bp downstream of exon 2 in *M. fascicularis*. Further investigation is required to be certain how to distinguish *HBA1* and *HBA2* in *M. mulatta*. The Illumina Hiseq reads used in the *M. mulatta* whole genome sequencing were 100 bp long, which means that it is not possible to distinguish *HBA1* and *HBA2* reads. A genotype such as GCY (GCC/GCT) at amino acid position 69 could therefore indicate (i) heterozygosity at *HBA1*, *HBA2* or both, or (ii) that animal encodes amino acid 69 with GCC in *HBA1* and GCT in *HBA2*, or vice versa.

#### 2. Supplementary Tables

**Supplementary table 1. Haemoglobin variation across the Primate Order.** Globin gene variation in this table includes copy number, electrophoretic, and amino acid variation. Lemur haemoglobin were not separated into alpha and beta chains, so variation recorded is not specific to either chain of haemoglobin.

| | Species | Alpha globin variation (number sampled) | Beta globin variation (number sampled) | References |
|---|---|---|---|---|
| Lemuroidea (lemurs) | *Lemur catta* | Y (28) | Y (28) | (Buettner-Janusch, et al. 1971) |
| | *Lemur fulvus* | N (63) | N (63) | (Buettner-Janusch, et al. 1971) |
| | *Lemur rufus* | N (29) | N (29) | (Buettner-Janusch, et al. 1971) |
| | *Propithecus verreauxi* | N (15) | N (15) | (Buettner-Janusch, et al. 1971) |
| Ceboidea (new world monkeys) | *Ateles belzebuth* | Y (6) | | (Boyer, et al. 1971) |
| | *Callicebus moloch* | | Y (27) | (Boyer, et al. 1971) |
| | *Cebus albifrons* | | Y (12) | (Boyer, et al. 1971) |
| | *Saguinus mystax* | | Y (56) | (Boyer, et al. 1971) |
| | *Saimiri sciureus* | | Y (7) | (Boyer, et al. 1971) |
| Cercopithecoidea (old world monkeys) | *Macaca* species | Y (6904) | Y (6904) | Supplementary tables 2 and 9 |
| | *Cercocebus* atys | Y (18) | N (18) | (Barnicot and Hewett-Emmett 1972) |
| | *Presbytis entellus* | N (26) | N (26) | (Hrdy, et al. 1975) |
| | *Presbytis cristatus* | N (21) | N (21) | (Ishimoto and Prychodko 1970) |
| Hominoidea (apes) | *Hylobates* species | N (22) | Y (23) | (Hoffman, et al. 1967; Boyer 1972; Zimmer, et al. 1980) |
| | *Pongo pygameaus* | Y (85) | | (Sullivan and Nute 1968; Takenaka, et al. 1993; Steiper, et al. 2006) |
| | *Pan troglodytes* | Y (151) | Y (151) | (Hoffman, et al. 1967; Boyer, et al. 1971; Zimmer, et al. 1980; Takenaka, et al. 1993) |
| | *Gorilla gorilla* | Y (5) | | (Boyer, et al. 1971; Boyer 1972; Zimmer, et al. 1980) |

**Supplementary table 2. Alpha globin phenotypic survey data.** Each row of the table represents a batch of long-tailed macaques for which a specific geographical origin is known. Numbers of long-tailed macaques exhibiting the electrophoretic phenotypes determined by the A, Q and P bands (see main text) are given for each batch. Only two studies (Barnicot, et al. (1966) and Ishimoto, et al. (1970)) reported the frequencies of the haemoglobin X band. For these studies, we have indicated in brackets the number of animals belonging to each of the A/Q/P phenotypes that also displayed a haemoglobin X band. For the Vietnam sample from Barnicot and colleagues (1966) we have added an extra row in grey italics indicating the distribution of haemoglobin X because only 122 of the 124 animals for which A/Q/P phenotypes were reported had the presence/absence of X reported. Only a subset of the samples in the table could be used in the malaria virulence analysis, owing to the patchiness of available malaria data. Samples used, and the spatial scale at which malaria virulence was determined, are indicated by the 'm' superscript; reasons for exclusion of other samples are indicated by numbered superscripts: [1]there was no corresponding macaque malaria data from the island or country, [2]the malaria data only reported presence of malaria and did not identify *Plasmodium* to species level, or [3] the reported origin of hemoglobin samples covered multiple islands with a range of possible levels of malaria virulence. However, scenarios involving these data are detailed in Supplementary figures 3 and 4.

| Location | | | | Phenotype | | | | | | | |
|---|---|---|---|---|---|---|---|---|---|---|---|
| Country | | Region | Total | A | AQ | Q | P | AP | QP | AQP | Refs |
| Cambodia[m] | | | 113 | 111 | 2 | 0 | 0 | 0 | 0 | 0 | (Ishimoto 1972) |
| Indonesia | | Bali[m] | 136 | 133 | 3 | 0 | 0 | 0 | 0 | 0 | (Kawamoto, et al. 1984) |
| Indonesia | | Bali[m] | 13 | 13 | 0 | 0 | 0 | 0 | 0 | 0 | (Kawamoto and Ischak 1981) |
| Indonesia | | Lombok[1] | 33 | 31 | 2 | 0 | 0 | 0 | 0 | 0 | (Kawamoto, et al. 1984) |
| Indonesia | Banten | Java[m] | 5 | 2 | 3 | 0 | 0 | 0 | 0 | 0 | (Kawamoto, et al. 1984) |
| Indonesia | Jawa Barat | Java[m] | 103 | 36 | 67 | 0 | 0 | 0 | 0 | 0 | (Kawamoto and Ischak 1981) |
| Indonesia | Jawa Barat | Java[m] | 137 | 73 | 64 | 0 | 0 | 0 | 0 | 0 | (Kawamoto, et al. 1984) |
| Indonesia | Jawa Barat | Java[m] | 88 | 53 | 35 | 0 | 0 | 0 | 0 | 0 | (Perwitasari-Farajallah, et al. 1999) |
| Indonesia | Jawa Tengah | Java[m] | 16 | 13 | 3 | 0 | 0 | 0 | 0 | 0 | (Kawamoto, et al. 1984) |
| Indonesia | Jawa Timur | Java[m] | 21 | 10 | 11 | 0 | 0 | 0 | 0 | 0 | (Kawamoto, et al. 1984) |
| Indonesia | Bengkulu | Sumatra[2] | 21 | 2 | 19 | 0 | 0 | 0 | 0 | 0 | (Kawamoto and Ischak 1981) |
| Indonesia | Lampung | Sumatra[2] | 70 | 16 | 54 | 0 | 0 | 0 | 0 | 0 | (Kawamoto and Ischak 1981) |
| Indonesia | Sumatera Barat | Sumatra[2] | 75 | 9 | 66 | 0 | 0 | 0 | 0 | 0 | (Kawamoto, et al. 1984) |
| Indonesia | Sumatera Selatan | Sumatra[2] | 13 | 4 | 9 | 0 | 0 | 0 | 0 | 0 | (Kawamoto and Ischak 1981) |
| Indonesia | Sumatera Utara | Sumatra[2] | 43 | 4 | 37 | 0 | 0 | 2 | 0 | 0 | (Kawamoto, et al. 1984) |
| Indonesia | | Sumbawa[1] | 78 | 0 | 78 | 0 | 0 | 0 | 0 | 0 | (Kawamoto, et al. 1984) |
| Indonesia[3] | | | 65 | 13 | 49 | 0 | 0 | 2 | 0 | 1 | (Smith and Ferrell 1980) |
| Indonesia | | | 10 | 9 | 1 | 0 | 0 | 0 | 0 | 0 | (Nozawa, et al. 1977) |
| Malaysia | | Peninsular[m] | 262 | 38 (12) | 190 (25) | 0 | 0 | 13 (3) | 21 (0) | 0 | (Ishimoto, et al. 1970) |
| Malaysia | | Peninsular[m] | 231 | 38 | 181 | 0 | 0 | 8 | 0 | 4 | (Barnicot, et al. 1970) |
| Malaysia | | Peninsular[m] | 184 | 41 | 129 | 0 | 1 | 2 | 0 | 11 | (Smith and Ferrell 1980) |
| Philippines | | Mindinao[2] | 59 | 0 | 59 (0) | 0 | 0 | 0 | 0 | 0 | (Ishimoto, et al. 1970) |
| Philippines[3] | | | 59 | 0 | 59 | 0 | 0 | 0 | 0 | 0 | (Ishimoto 1972) |
| Singapore[m] | | | 10 | 7 (0) | 0 | 0 | 1 (0) | 2 (2) | 0 | 0 | (Barnicot and Jolly 1966) |
| Thailand[m] | | | 186 | 91 (26) | 61 (19) | 0 | 2 (0) | 23 (5) | 9 (2) | 0 | (Ishimoto, et al. 1970) |
| Thailand[m] | | | 68 | 41 (12) | 14 (3) | 1 (0) | 1 (0) | 9 (2) | 2 (0) | 0 | (Barnicot and Jolly 1966) |
| Thailand[m] | South | | 33 | 2 | 26 | 0 | 0 | 5 | 0 | 0 | (Kawamoto, et al. 1989) |
| Thailand[m] | Central | | 245 | 244 | 1 | 0 | 0 | 0 | 0 | 0 | (Kawamoto, et al. 1989) |
| Vietnam[1] | | | 124 | 77 | 34 | 2 | 1 | 8 | 2 | 0 | (Barnicot and Jolly 1966) |
| *Vietnam[1]* | | | *122* | *76 (11)* | *33 (10)* | *2 (0)* | *1 (0)* | *8 (2)* | *2 (0)* | *0* | *(Barnicot and Jolly 1966)* |

Supplementary table 2 (legend on previous page)

**Supplementary table 3. The prevalence of *Plasmodium* species infection among *M. fascicularis*.** "Presence of virulent species" refers to at least one of *P. knowlesi* and *P. coatneyi* being present. All historical surveys (pre-1990) used microscopy and morphological traits to identify *Plasmodium* species from blood. Data obtained using morphological methods as opposed to molecular methods are listed separately because the use of PCR greatly increases the probability of detecting parasites (Putaporntip, et al. 2010). We also searched the literature for case reports to supplement surveys; Coatney, et al. (1971) confirm presence of *P. knowlesi*, *P. coatneyi* in Cebu and Palawan (both in the Philippines).

| **Country** | **location** | **number of surveys** (morphological/ molecular) | **number of individuals** (morphological/ molecular) | **total infected** (morphological/ molecular) | **malaria species richness** | **presence of virulent species** | **refs** |
|---|---|---|---|---|---|---|---|
| Cambodia | Not specified | 2/0 | 51/0 | 13/0 | 1 | N | (Fooden 1994)* |
| Thailand | Not specified | 1/6 | 8/307 | 2/20 | 4 | Y | (Seethamchai, et al. 2008; Putaporntip, et al. 2010) |
| Malaysia | Peninsular | 14/4 | 477/235 | 152/144 | 5 | Y | (Fooden 1994; Vythilingam, et al. 2006; Ho, et al. 2010) |
| | Sarawak | 0/1 | 0/82 | 0/80 | 5 | Y | (Lee, et al. 2011) |
| | Not specified | 5/0 | 1110/0 | 256/0 | 4 | Y | (Fooden 1994) |
| Singapore | | 0/1 | 0/13 | 0/3 | 1 | Y | (Jeslyn, et al. 2011) |
| Indonesia | Sumatra | 1/0 | 235/0 | 20/0 | unknown ^ | - | (Fooden 1994) |
| | Java | 5/0 | 114/0 | 10/0 | 2 | Y | (Fooden 1994) |
| | Bali | 1/0 | 60/0 | 0/0 | 0 | N | (Fooden 1994) |
| | Not specified | 1/0 | 354/0 | 50 | 1 | N | (Fooden 1994) |
| Philippines | Palawan | 1/0 | 20/0 | 11/0 | 2 | Y | (Fooden 1994) |
| | Palawan & Mindanao | 1/0 | 162/0 | 16/0 | unknown ^ | - | (Fooden 1994) |
| | Cebu | 2/0 | 174/0 | 12/0 | 2 | Y | (Fooden 1994) |
| | Not specified | 2/0 | 181/0 | 25/0 | 1 | N | (Fooden 1994) |

^ All positive malaria samples were of unknown malaria species (Howard and Cabrera 1961; Matsubayashi and Sajuthi 1981); other authors also reported unknown malaria species but in areas where there were several other surveys.* Primary references detailed in Fooden (1994).

**Supplementary table 4. Heterozygosity of *Macaca fascicularis* populations from different locations at non-globin loci.** Here we summarise the observed heterozygosity of *M. fascicularis* from a range of geographical locations at 9 different types of non-globin loci. Wherever possible we indicate the sample number (N), but in some sources this was not stated. Estimates for named loci (e.g. Haptoglobin) in studies from the 1960s-80s may sometimes be dervied from exactly the same macaque samples used for the relevant populations in supplementary table 2, but this is hard to fully ascertain. More modern studies report heterozygosity among macaques from different geographical origins using panels of short tandem repeat (STR) or single nucleotide polymorphism (SNP) markers. We have included such estimates from a recent study of relevant geographical populations (Day, et al. 2018). [$]A possible haptoglobin deficiency was observed by Kawamoto and Ischak in one macaque from Sumatra, but they could not be certain if this was a genetically determined deficiency. [#]Acp and PHI loci were reported to have been examined by Kawamoto and Ischak (1981), but no polymorphism was reported in the results, hence we assume zero heterozygosity, and that the number of animals examined was similar to the numbers examined for the other loci.

| **Location** | **Transferrin (Tf)** | **Haptoglobin (Hp)** | **Albumin (Alb)** | **Acid phosphatase (AP or Acp)** | **6 phosphoglucanate dehydrogenase (PGD)** | **Phosphohexose isomerase (PHI)** | **NADH-dependent diaphorase (Dia)** | **STR markers** | **SNP markers** | **References** |
|---|---|---|---|---|---|---|---|---|---|---|
| **Philippines (unspecified)** | 0.02 | 0 | 0 | 0 | 0 | 0.49 | 0 | - | - | (Ishimoto 1973) |
| **Philippines (Mindanao)** | - | - | - | - | - | - | - | 0.64 (N= 37) | 0.1 (N=20) | (Day, et al. 2018) |
| **Thailand** | 0.72 | 0 | 0 | 0.17 | 0.07 | 0.08 | 0.43 | - | - | (Ishimoto 1973) |
| **Malaysia** | 0.53 | 0 | 0 | 0.03 | 0.24 | 0.02 | 0.06 | - | - | (Ishimoto 1973) |
| **Cambodia** | 0.78 (N=114) | - | - | 0.07 (N=113) | - | - | - | 0.72 (N=103) | 0.12 (N= 23) | (Ishimoto, et al. 1968; Toyomasu and Ishimoto 1969; Day, et al. 2018) |
| **Sumatra** | 0.53 (N=103) | 0[$] (N=104) | 0.029 (N=104) | 0[#] | 0.06 (N=100) | 0[#] | 0.030 (N= 99) | 0.71 (N= 97) | 0.14 (N= 38) | (Kawamoto and Ischak 1981; Day, et al. 2018) |
| **Java** | 0.57 (N=105) | 0 (N=110) | 0 (N=109) | 0[#] | 0.02 (N=100) | 0[#] | 0.020 (N= 98) | - | - | (Kawamoto and Ischak 1981) |
| **Bali** | 0 (N=13) | 0 (N=13) | 0 (N=13) | 0[#] (N=13) | 0 (N=13) | 0[#] (N=13) | 0 (N= 13) | - | - | (Kawamoto and Ischak 1981) |
| **Singapore** | - | - | - | - | - | - | - | 0.66 (N= 60) | 0.15 (N= 25) | (Day, et al. 2018) |

**Supplementary table 5. Primers for amplifying *M. fascicularis HBA1* and *HBA2*.** Primer sequences and annealing conditions to amplify exon 2 within a nested PCR reaction that first amplifies *HBA1* and *HBA2* separately.

| **name** | **forward (F) /reverse (R)** | **sequence (5' to 3')** | **annealing temp** | **target region** | **product length** |
|---|---|---|---|---|---|
| F3F | F | TCTGGTCCCCACAGACTCAG | 70°C | *HBA1* | 820 |
| F8R | R | CTGCAGAGAGGTCCTAGCCA | | | |
| F3F | F | TCTGGTCCCCACAGACTCAG | 68°C | *HBA2* | 820 |
| F4R | R | GAGAGAGAGAACCCAGGCAC | | | |
| MM2 | F | CCACCCCTCACTCTGCTTCT | 68°C | exon 2 (both *HBA1* & *HBA2*) | 374 |
| S3R | R | GTGCAGAGAAGAGGGTCAGT | | | |

**Supplementary table 6. Proposed *M. fascicularis* alpha globin haplotypes present in our sample.** As noted in the main text, it is not possible to unequivocally phase the alpha globin haplotypes present in our sample. However, we manually ascertained a possible set of haplotypes that could account for the patterns seen, on the assumption that each alpha globin cluster contains exactly one copy of *HBA1* and up to three copies of *HBA2*. Here we display 23 possible haplotypes, which between them can account for the genotypes of 70 of the 77 animals for which we obtained alpha globin exon 2 sequences. The order of alpha globin genes indicated in each haplotype is arbitrary. We also note that a haplotype which is indicated here to contain a single copy of an *HBA1* or *HBA2* sequence might in fact contain repeated copies of that same sequence. We have coloured each alpha globin sequence name by the amino acids it encodes at variable positions, also described in figure 2 of the main text. The *HBA1* amino acid sequences are: HBA1_GTGVH, HBA1_DTGVH, HBA1_GIRRQ, HBA1_GTGVQ and HBA1_GTRVQ. The *HBA2* amino acid sequences are: HBA2_VEH, HBA2_VGQ, HBA2_VGH and HBA2_IGH. If only amino acid differences and numbers of genes are considered (i.e. synonymous differences between alpha globin sequences are ignored), there are 16 different haplotypes in this proposed scheme.

| Haplotype number | Predicted sequences in haplotype | Frequency in sample |
|---|---|---|
| **1** | HBA1.1-HBA2.3 | 1 |
| **2** | HBA1.1-HBA2.5 | 12 |
| **3** | HBA1.1-HBA2.6 | 7 |
| **4** | HBA1.1-HBA2.8 | 6 |
| **5** | HBA1.1-HBA2.1-HBA2.2 | 22 |
| **6** | HBA1.2-HBA2.3 | 18 |
| **7** | HBA1.2-HBA2.5 | 5 |
| **8** | HBA1.2-HBA2.1-HBA2.2-HBA2.5 | 3 |
| **9** | HBA1.3-HBA2.4-HBA2.7 | 3 |
| **10** | HBA1.3-HBA2.1-HBA2.2-HBA2.4 | 11 |
| **11** | HBA1.4-HBA2.3 | 8 |
| **12** | HBA1.4-HBA2.2-HBA2.7 | 3 |
| **13** | HBA1.4-HBA2.9-HBA2.10 | 2 |
| **14** | HBA1.5-HBA2.1-HBA2.2 | 4 |
| **15** | HBA1.5-HBA2.1-HBA2.2-HBA2.4 | 7 |
| **16** | HBA1.6-HBA2.1 | 5 |
| **17** | HBA1.6-HBA2.1-HBA2.2 | 3 |
| **18** | HBA1.7-HBA2.1-HBA2.2 | 5 |
| **19** | HBA1.8-HBA2.2 | 2 |
| **20** | HBA1.8-HBA2.7 | 3 |
| **21** | HBA1.9-HBA2.1-HBA2.2 | 4 |
| **22** | HBA1.10-HBA2.1-HBA2.2-HBA2.4 | 3 |
| **23** | HBA1.11-HBA2.1-HBA2.2-HBA2.4 | 3 |

**Supplementary table 7. Haplotype assignments and parental or sibling relationships within our sample.** Each row of the table refers to an animal within our sample. The numbers in the "predicted haplotype combination" column refer to the predicted haplotypes in Supplementary table 6. These predictions were made manually on the assumption that each alpha globin cluster contains exactly one copy of HBA1 and up to three copies of HBA2. We only predicted haplotypes if we required that haplotype to exist in at least 2 different animals, with the exception of haplotype 1, for which there is reasonable evidence based on animal Mf1_59 (although it remains possible that Mf1_59 is a homozygote for "HBA1.1-HBA1.4-HBA2.3"). In this way, we were able to predict possible haplotypes for 70 of the 77 animals in our sample. Some of the remaining 7 animals look as though they may possess some of our predicted haplotypes, plus an as yet unpredicted haplotype. However, we have not made predictions for those animals on that basis, since this would involve creating "singleton" predicted haplotypes, with little evidence to support them. There are familial relationships between some of the animals in our sample, and we checked whether our haplotype predictions are consistent with these known familial relationships. Maternal/offspring relationships are reliably known, but paternal relationships are harder to obtain for all these samples. Three potential paternal relationships are indicated. Since paternity is uncertain, "sibling" in our table indicates animals which share a mother, but may only be half siblings. Our haplotype predictions are consistent with all familial relationships, with the exception that Mf1_77 is the mother of Mf1_88. Mf1_77 seems to possess only HBA1.1 as its *HBA1* sequence and Mf1_88 seems to possess only HBA1.8 as its *HBA1* sequence, which makes it impossible to define a haplotype they both share. However, as noted in the methods, we used a 2% cutoff to determine the presence of alpha globin sequences. If we examine all sequences from Mf1_88, we see that it does have HBA1.1, at a frequency below this threshold (1.63% of reads; n =61/3731). It is likely that Mf1_77 and Mf1_88 share the HBA1.1 sequence (and might share haplotype "5" within our scheme), and there was unequal amplification of HBA1.1 in Mf1_88. The read depth of *HBA1* in Mf1_88 is low (3731 reads) compared to the mean read depth of *HBA1* for an individual across all our samples (9393.65 reads).

| SampleID | Predicted haplotypes | Sequences present in sample | Parental or sibling relationships |
|---|---|---|---|
| 'Mf1_13' | [7,18] | ',HBA1.2,HBA1.7,HBA2.1,HBA2.2,HBA2.5' | mother of Mf1_14 |
| 'Mf1_14' | [6,18] | ',HBA1.2,HBA1.7,HBA2.1,HBA2.2,HBA2.3' | offspring of Mf1_13 |
| 'Mf1_15' | [7,20] | ',HBA1.2,HBA1.8,HBA2.5,HBA2.7' | |
| 'Mf1_16' | [5,18] | ',HBA1.1,HBA1.7,HBA2.1,HBA2.2' | |
| 'Mf1_17' | [2,14] | ',HBA1.1,HBA1.5,HBA2.1,HBA2.2,HBA2.5' | mother of Mf1_27 |
| 'Mf1_18' | [5,11] | ',HBA1.1,HBA1.4,HBA2.1,HBA2.2,HBA2.3' | sibling of Mf1_62 |
| 'Mf1_19' | ? | ',HBA1.1,HBA1.3,HBA2.1,HBA2.2,HBA2.5,HBA2.12' | |
| 'Mf1_20' | [2,8] | ',HBA1.1,HBA1.2,HBA2.1,HBA2.2,HBA2.5' | |
| 'Mf1_21' | [6,17] | ',HBA1.2,HBA1.6,HBA2.1,HBA2.2,HBA2.3' | |
| 'Mf1_22' | [6,13] | ',HBA1.2,HBA1.4,HBA2.3,HBA2.9,HBA2.10' | |
| 'Mf1_23' | ? | ',HBA1.2,HBA1.13,HBA2.1,HBA2.2,HBA2.3' | |
| 'Mf1_24' | [16,19] | ',HBA1.6,HBA1.8,HBA2.1,HBA2.2' | |
| 'Mf1_25' | [5,16] | ',HBA1.1,HBA1.6,HBA2.1,HBA2.2' | |
| 'Mf1_26' | ? | ',HBA1.1,HBA1.2,HBA2.3,HBA2.5,HBA2.6' | sibling of Mf1_52 |
| 'Mf1_27' | [2,20] | ',HBA1.1,HBA1.8,HBA2.5,HBA2.7' | offspring of Mf1_17 |
| 'Mf1_28' | [6,7] | ',HBA1.2,HBA2.3,HBA2.5' | |
| 'Mf1_29' | [10,10] | ',HBA1.3,HBA2.1,HBA2.2,HBA2.4' | |
| 'Mf1_30' | [4,22] | ',HBA1.1,HBA1.10,HBA2.1,HBA2.2,HBA2.4,HBA2.8' | |
| 'Mf1_31' | [5,11] | ',HBA1.1,HBA1.4,HBA2.1,HBA2.2,HBA2.3' | |
| 'Mf1_32' | [3,5] | ',HBA1.1,HBA2.1,HBA2.2,HBA2.6' | offspring of Mf1_75 and possibly Mf1_72, may therefore share a father with 87 |
| 'Mf1_33' | [10,18] | ',HBA1.3,HBA1.7,HBA2.1,HBA2.2,HBA2.4' | |
| 'Mf1_34' | ? | ',HBA1.1,HBA1.4,HBA2.3,HBA2.9,HBA2.11' | |
| 'Mf1_35' | [5,6] | ',HBA1.1,HBA1.2,HBA2.1,HBA2.2,HBA2.3' | |
| 'Mf1_36' | [6,6] | ',HBA1.2,HBA2.3' | |
| 'Mf1_37' | ? | ',HBA1.2,HBA1.3,HBA2.3,HBA2.4,HBA2.10,HBA2.11' | |
| 'Mf1_38' | [3,6] | ',HBA1.1,HBA1.2,HBA2.3,HBA2.6' | |
| 'Mf1_39' | [2,18] | ',HBA1.1,HBA1.7,HBA2.1,HBA2.2,HBA2.5' | |

| | | | |
|---|---|---|---|
| 'Mf1_40' | [2,5] | ',HBA1.1,HBA2.1,HBA2.2,HBA2.5' | |
| 'Mf1_41' | [5,10] | ',HBA1.1,HBA1.3,HBA2.1,HBA2.2,HBA2.4' | |
| 'Mf1_42' | [5,10] | ',HBA1.1,HBA1.3,HBA2.1,HBA2.2,HBA2.4' | |
| 'Mf1_43' | [5,6] | ',HBA1.1,HBA1.2,HBA2.1,HBA2.2,HBA2.3' | |
| 'Mf1_44' | [11,15] | ',HBA1.4,HBA1.5,HBA2.1,HBA2.2,HBA2.3,HBA2.4' | sibling of Mf1_60 |
| 'Mf1_45' | [6,17] | ',HBA1.2,HBA1.6,HBA2.1,HBA2.2,HBA2.3' | |
| 'Mf1_46' | [12,21] | ',HBA1.4,HBA1.9,HBA2.1,HBA2.2,HBA2.7' | |
| 'Mf1_47' | [5,16] | ',HBA1.1,HBA1.6,HBA2.1,HBA2.2' | mother of Mf1_63, sibling of Mf1_56 |
| 'Mf1_48' | [2,15] | ',HBA1.1,HBA1.5,HBA2.1,HBA2.2,HBA2.4,HBA2.5' | |
| 'Mf1_49' | [2,8] | ',HBA1.1,HBA1.2,HBA2.1,HBA2.2,HBA2.5' | |
| 'Mf1_50' | [10,11] | ',HBA1.3,HBA1.4,HBA2.1,HBA2.2,HBA2.3,HBA2.4' | |
| 'Mf1_52' | [4,6] | ',HBA1.1,HBA1.2,HBA2.3,HBA2.8' | sibling of Mf1_26 |
| 'Mf1_53' | [10,13] | ',HBA1.3,HBA1.4,HBA2.1,HBA2.2,HBA2.4,HBA2.9,HBA2.10' | |
| 'Mf1_54' | [2,5] | ',HBA1.1,HBA2.1,HBA2.2,HBA2.5' | |
| 'Mf1_55' | [6,6] | ',HBA1.2,HBA2.3' | |
| 'Mf1_56' | [5,16] | ',HBA1.1,HBA1.6,HBA2.1,HBA2.2' | |
| 'Mf1_57' | [2,21] | ',HBA1.1,HBA1.9,HBA2.1,HBA2.2,HBA2.5' | |
| 'Mf1_58' | [6,15] | ',HBA1.2,HBA1.5,HBA2.1,HBA2.2,HBA2.3,HBA2.4' | |
| 'Mf1_59' | [1,11] | ',HBA1.1,HBA1.4,HBA2.3' | |
| 'Mf1_60' | [2,15] | ',HBA1.1,HBA1.5,HBA2.1,HBA2.2,HBA2.4,HBA2.5' | sibling of Mf1_44 |
| 'Mf1_61' | [11,17] | ',HBA1.4,HBA1.6,HBA2.1,HBA2.2,HBA2.3' | |
| 'Mf1_62' | [5,11] | ',HBA1.1,HBA1.4,HBA2.1,HBA2.2,HBA2.3' | sibling of Mf1_18 |
| 'Mf1_63' | [16,21] | ',HBA1.6,HBA1.9,HBA2.1,HBA2.2' | offspring of Mf1_47 |
| 'Mf1_64' | [3,6] | ',HBA1.1,HBA1.2,HBA2.3,HBA2.6' | |
| 'Mf1_65' | [5,15] | ',HBA1.1,HBA1.5,HBA2.1,HBA2.2,HBA2.4' | |
| 'Mf1_66' | ? | ',HBA1.3,HBA1.12,HBA2.1,HBA2.2,HBA2.4,HBA2.7' | |
| 'Mf1_67' | [10,20] | ',HBA1.3,HBA1.8,HBA2.1,HBA2.2,HBA2.4,HBA2.7' | mother of Mf1_71 |
| 'Mf1_68' | [2,15] | ',HBA1.1,HBA1.5,HBA2.1,HBA2.2,HBA2.4,HBA2.5' | |
| 'Mf1_69' | [7,10] | ',HBA1.2,HBA1.3,HBA2.1,HBA2.2,HBA2.4,HBA2.5' | |
| 'Mf1_70' | [11,14] | ',HBA1.4,HBA1.5,HBA2.1,HBA2.2,HBA2.3' | |
| 'Mf1_71' | [10,14] | ',HBA1.3,HBA1.5,HBA2.1,HBA2.2,HBA2.4' | offspring of Mf1_67 |
| 'Mf1_72' | [5,19] | ',HBA1.1,HBA1.8,HBA2.1,HBA2.2' | father of Mf1_90, possible father of Mf1_87 and Mf1_32, sibling of Mf1_86 |
| 'Mf1_73' | [4,9] | ',HBA1.1,HBA1.3,HBA2.4,HBA2.7,HBA2.8' | |
| 'Mf1_74' | [6,10] | ',HBA1.2,HBA1.3,HBA2.1,HBA2.2,HBA2.3,HBA2.4' | |
| 'Mf1_75' | [3,4] | ',HBA1.1,HBA2.6,HBA2.8' | mother of Mf1_32 |
| 'Mf1_76' | [2,4] | ',HBA1.1,HBA2.5,HBA2.8' | |
| 'Mf1_77' | [4,5] | ',HBA1.1,HBA2.1,HBA2.2,HBA2.8' | mother of Mf1_88 |
| 'Mf1_78' | [14,23] | ',HBA1.5,HBA1.11,HBA2.1,HBA2.2,HBA2.4' | |
| 'Mf1_79' | [3,8] | ',HBA1.1,HBA1.2,HBA2.1,HBA2.2,HBA2.5,HBA2.6' | |
| 'Mf1_80' | [12,21] | ',HBA1.4,HBA1.9,HBA2.1,HBA2.2,HBA2.7' | |
| 'Mf1_81' | [6,23] | ',HBA1.2,HBA1.11,HBA2.1,HBA2.2,HBA2.3,HBA2.4' | |
| 'Mf1_82' | [3,5] | ',HBA1.1,HBA2.1,HBA2.2,HBA2.6' | |
| 'Mf1_83' | [6,23] | ',HBA1.2,HBA1.11,HBA2.1,HBA2.2,HBA2.3,HBA2.4' | |
| 'Mf1_84' | [3,22] | ',HBA1.1,HBA1.10,HBA2.1,HBA2.2,HBA2.4,HBA2.6' | mother of Mf1_87 |
| 'Mf1_85' | [7,9] | ',HBA1.2,HBA1.3,HBA2.4,HBA2.5,HBA2.7' | |
| 'Mf1_86' | [5,9] | ',HBA1.1,HBA1.3,HBA2.1,HBA2.2,HBA2.4,HBA2.7' | sibling of Mf1_72 |
| 'Mf1_87' | [5,22] | ',HBA1.1,HBA1.10,HBA2.1,HBA2.2,HBA2.4' | offspring of Mf1_84 and possibly Mf1_72 |
| 'Mf1_88' | ? | ',HBA1.8,HBA2.1,HBA2.2,HBA2.8' | offspring of Mf1_77 |
| 'Mf1_89' | [5,15] | ',HBA1.1,HBA1.5,HBA2.1,HBA2.2,HBA2.4' | |
| 'Mf1_90' | [5,12] | ',HBA1.1,HBA1.4,HBA2.1,HBA2.2,HBA2.7' | offspring of Mf1_72 |

**Supplementary table 8. Summary of *Macaca mulatta* exon 2 sequences for sites which are variable.** This table summarises the strength of evidence for variation, or lack of variation, in specific sites in *M. mulatta* alpha globin exon 2. See section 1.5 for further details. Supplementary table 8 includes all the exon 2 sites found to vary in *M. fascicularis* (Main text) and the two sites that we found to be variable in *M. mulatta* (see section 1.1.5). The number of *M. mulatta* individuals for which we have data varies from site to site owing to differences in the quality of data for different animals. We use ambiguity codes as follows: A or T = W; A or G = R; A or C = M; T or G = K; T or C = Y; C or G = S.

| **Amino acid position in complete alpha globin sequence.** | **Nucleotide acid position (start of codon) in exon 2** | ***M. mulatta* results** | ***M. mulatta* amino acid** |
|---|---|---|---|
| 44 | 37 | 91/91 CCC | Pro |
| 52 | 61 | 86/86 TCT | Ser |
| 55 | 70 | 87/87 GTT | Val |
| 57 | 76 | 86/86 GGC | Gly |
| 59 | 82 | 83/84 GGC *(1/84 GRC, presumed error, read depth for site = 34)* | Gly |
| 66 | 103 | 70/71 CTG *(1/71 YTG, presumed error, read depth for site = 10)* | Leu |
| 67 | 106 | 69/69 ACC | Thr |
| 69 | 112 | 56/66 GCC; 10/66 GCY | Ala |
| 71 | 118 | 65/65 GGG | Gly |
| 73 | 124 | 64/64 GTG | Val |
| 78 | 139 | 42/55 CAC; 11/55 CAM; 2/55 CAA | His / Gln |

**Supplementary table 9. Haemoglobin surveys of the *Macaca* genus.** For the population based surveys (all but *M. assamensis* and *M. sinica*), we define the AQ phenotype as a report of both the wild type haemoglobin band (**A**) and a 'quicker' (**Q**), or more negative, band in gel electrophoresis.*Takenaka, et al. (1988) reference studies by Maita and colleagues that report a Glycine >Aspartic Acid amino acid polymorphism at position 15 in *M. assamensis* (exactly the same substitution as seems to cause an AQ phenotype in *M. arctoides*), and an Alanine > Aspartic Acid polymorphism at position 12 of alpha globin in *M. sinica*. However we were unable to find either Maita reference and do not know the original sample size, nor the exact electrophoretic behavior of these variants. Full publication details for these references are: Maita T, Tanioka Y, Shotake T, Matsuda G (1984) Adult hemoglobins from two major components from toque monkey (*Macaca sinica*). Seikagaku. 56(8):813. & Maita T, Yajima E,Komine H, Ushimi K, Matsuda G (1984) Amino acid sequences of three major components of adult hemoglobins from Assam's monkey (*Macaca assamenses*). Seikagaku. 56(8):813. They reference the same page, so may be conference abstracts, but as the variant haemoglobins are described in Takenaka *et al* (1988) in Table 1 in detail, we include them here. These two species are indicated in red in Figure 4, main text, but have a dotted line as we could not locate the original sample sizes.

| ***Macaca* species** | **individuals surveyed** | **number of publications** | **observed 'AQ'** | **observed any variation** | **references** |
|---|---|---|---|---|---|
| ***M. arctoides*** | 256 | 3 | Y | | (Kitchen, et al. 1968; Ishimoto, et al. 1970; Nozawa, et al. 1977) |
| ***M. assamensis*** | * | * | Likely* | Y | *(Takenaka, et al. 1988) |
| ***M. brunnescens*** | 17 | 1 | N | | (Takenaka, et al. 1987) |
| ***M. cyclopis*** | 301 | 3 | N | | (Ishimoto, et al. 1970; Ishimoto 1972; Nozawa, et al. 1977) |
| ***M. fascicularis*** | 2371 | 8 | Y | | Supplementary table 2 |
| ***M. fuscata*** | 2539 | 5 | N | Y – 22 (X) | (Ishimoto, et al. 1970; Ishimoto 1972; Nozawa, et al. 1977) |
| ***M. hecki*** | 34 | 1 | N | | (Takenaka, et al. 1987) |
| ***M. maura*** | 66 | 2 | N | | (Weiss, et al. 1973; Takenaka, et al. 1987) |
| ***M. mulatta*** | 633 | 4 | N | | (Ishimoto, et al. 1970; Ishimoto 1972; Nozawa, et al. 1977; Tomiuk 1989) |
| ***M. nemestrina*** | 510 | 3 | Y | | (Ishimoto, et al. 1970; Ishimoto 1972; Nozawa, et al. 1977) |
| ***M. nigra*** | 53 | 2 | N | | (Weiss, et al. 1973; Takenaka, et al. 1987) |
| ***M. nigrescens*** | 19 | 1 | N | | (Takenaka, et al. 1987) |
| ***M. ochreata*** | 18 | 1 | N | | (Takenaka, et al. 1987) |
| ***M. radiata*** | 38 | 2 | Y | | (Weiss, et al. 1973; Nozawa, et al. 1977) |
| ***M. sinica*** | * | * | Unknown* | Y | *(Takenaka, et al. 1988) |
| ***M. tonkeana*** | 49 | 1 | N | | (Takenaka, et al. 1987) |

**Supplementary table 10. Occurrence of *Plasmodium* species in *Macaca*.** The macaque genus is infected with seven malaria parasites: *P. coatneyi, P. cynomolgi, P. fieldi, P. fragile, P. knowlesi, P.inui,* and *P. simiovale* (Garnham 1966; Coatney, et al. 1971). Two (*P. fragile* and *P. simiovale*) are restricted to the Indian sub-continent and three (*P. coatneyi, P. fieldi,* and *P. knowlesi*) are restricted to Southeast Asia. *P. inui* and *P. cynomolgi* have been recorded from both regions in tropical broadleaf evergreen forests where the *Anopheles leucosphyrus* group of mosquitoes is found (Colless 1956). No parasite surveys have been published on the Barbary macaque, *M. sylvanus*, in Europe. *If there was more than three references for a given species, only three were listed in the table.

| | *P. coatneyi* | *P,cynomolgi* | *P. fieldi* | *P. fragile* | *P. inui* | *P. knowlesi* | *P. simiovale* | individuals surveyed | number of publications | references* |
|---|---|---|---|---|---|---|---|---|---|---|
| *M. arctoides* | ✓ | ✓ | ✓ | | ✓ | ✓ | | 29 | 2 | (Putaporntip, et al. 2010; Fungfuang, et al. 2020) |
| *M. cyclopis* | | ✓ | | | ✓ | | | 959 | 4 | (Chuang, et al. 1966; Peyton and Harrison 1980; Huang, et al. 2010) |
| *M. fascicularis* | ✓ | ✓ | ✓ | | ✓ | ✓ | | 3474 | 37 | See Supplementary table 3 |
| *M. fuscata* | | | | | | | | 547 | 1 | (Otsuru and Sekikawa 1979) |
| *M. mulatta* | | | | | ✓ | | | 24000+ | 7 | (Schmidt, et al. 1961; Prakash and Chakrabarti 1962) |
| *M. nemestrina* | ✓ | ✓ | ✓ | | ✓ | ✓ | | 636 | 6 | (Eyles, et al. 1962; Vythilingam, et al. 2008; Putaporntip, et al. 2010) |
| *M. nigra* | | | | | ✓ | | | 3 | 1 | (Eyles and Warren 1962) |
| *M. radiata* | | ✓ | | ✓ | ✓ | | | 318 | 6 | (Mulligan and Swaminath 1940; Ramakrishnan and Mohan 1962; Choudhury, et al. 1963) |
| *M. sinica* | | ✓ | | ✓ | ✓ | | ✓ | 47 | 3 | (Dissanaike 1963, 1965; Dissanaike, et al. 1965; Nelson 1971) |

#### 3. Supplementary Figures S1-S9

**Supplementary Figure 1. Malaria surveys in long-tailed macaques.** A. Map of malaria surveys in long-tailed macaques with corresponding haemoglobin phenotype surveys. B. A mosaic plot of cumulative number of *M. fascicularis* checked for malaria infections. The height of each block refers to the proportion of long-tailed macaques with a given infection status from within the country. Dark red highlights the proportion of long-tailed macaques with *Plasmodium knowlesi* or *P. coatneyi* infections. The width of each column is proportional to the number of individuals examined for malaria across Southeast Asia.

**A.**

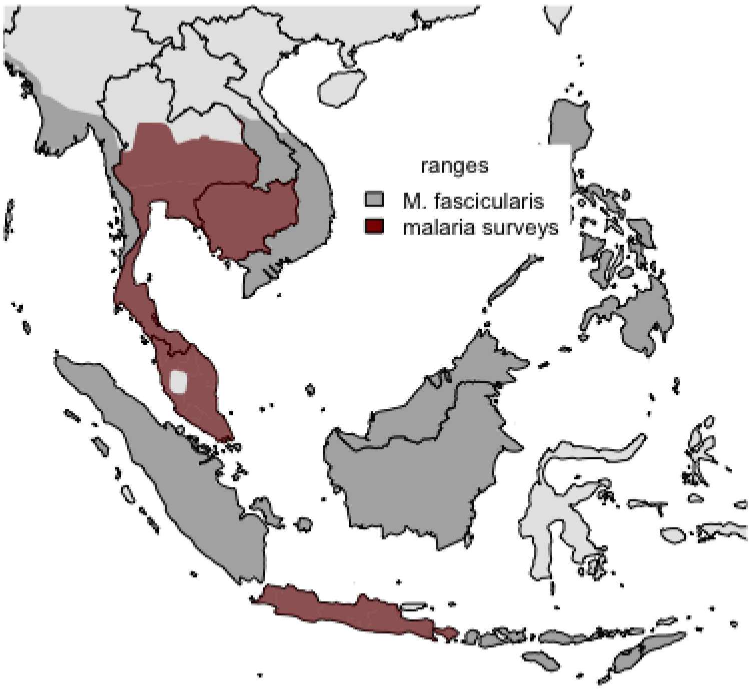


**B.**

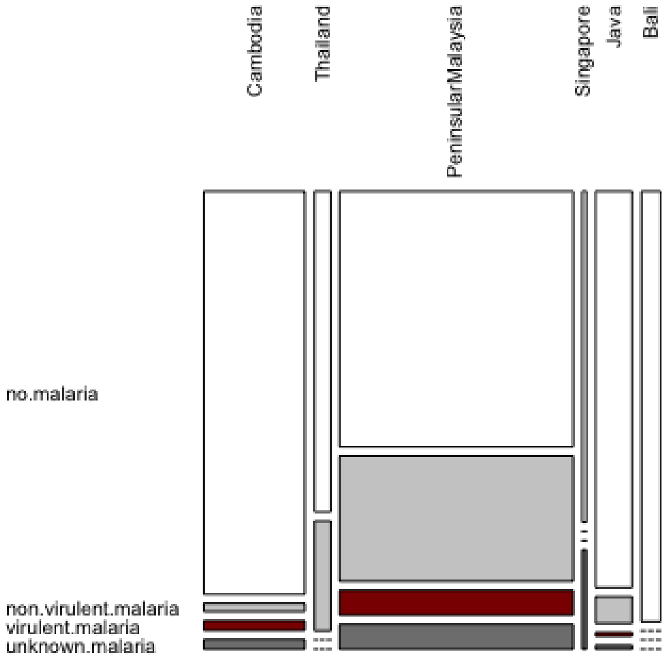

**Supplementary Figure 2. Robustness of the virulence analysis.** Increasing the cutoff for likelihood of non- virulent malarias did not have a large effect on the probability of observing a macaque with a variant phenotype. The histograms display the number of times the Metropolis Hastings sampler estimated a particular value (out of 1000 estimates).

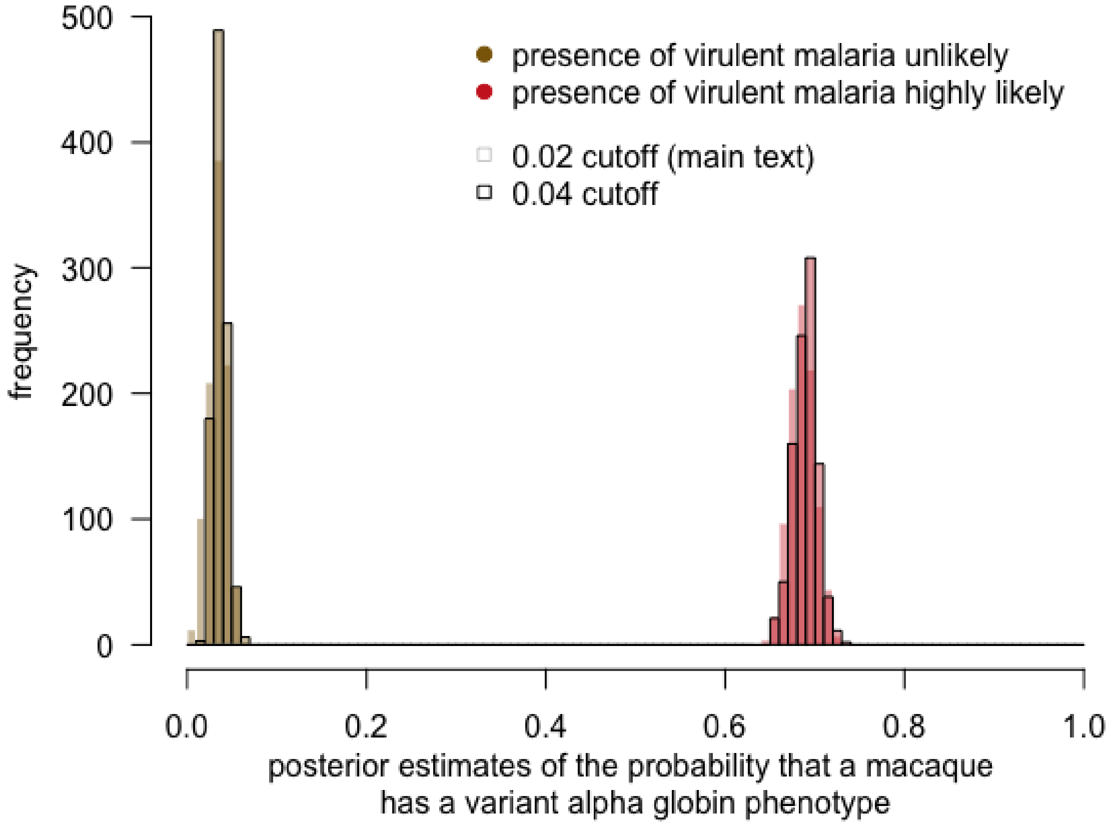

**Supplementary Figure 3. Varying haemoglobin surveys included in the analysis. (A)** Expands coverage to include Mindanao and Sumatran samples – either assuming the malarias found are non virulent or virulent. **(B)** We can also include all haemoglobin surveys by using malaria likelihoods from the malaria data from the closest survey. Neither of these change the posterior probability substantially.

**A.**

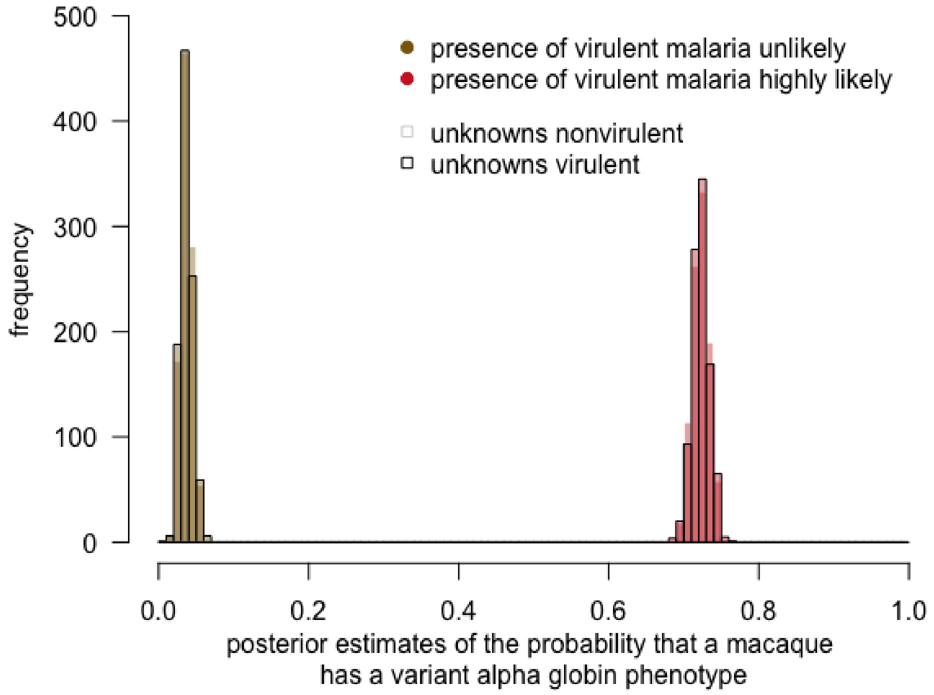


**B.**

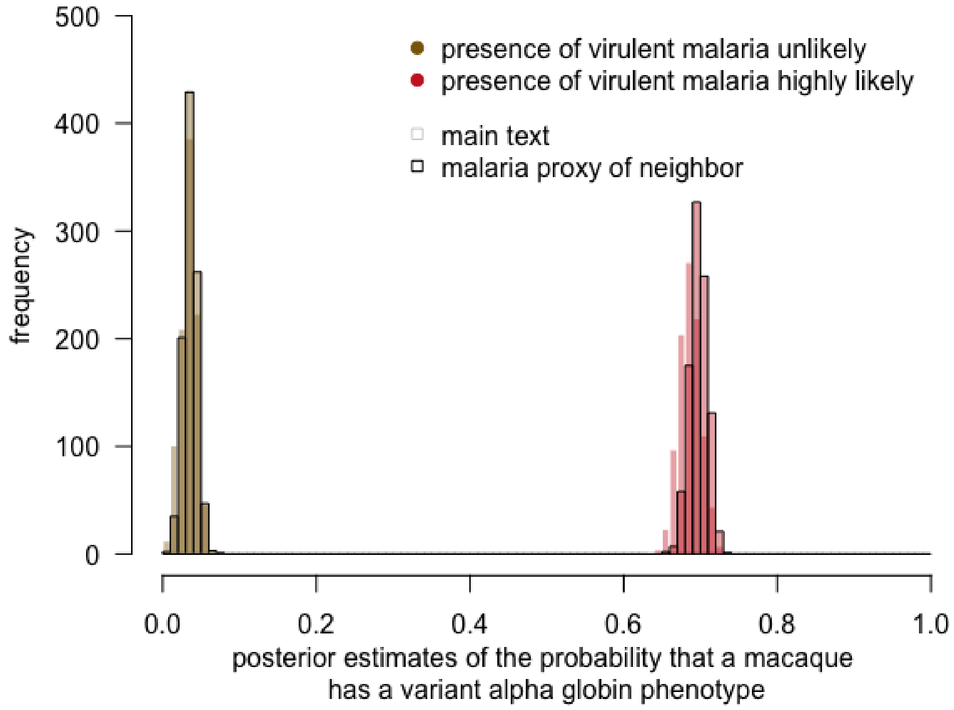

**Supplementary Figure 4. Non-A phenotype proportions in regions where minor haemoglobins were tested.** Only certain studies used protocols that allowed the detection of minor haemoglobins. If only major haemoglobins were included, it is much more likely to observe variants if virulent malarias are likely present. If minor haemoglobins are also considered variants, then most long-tailed macaques have variant haemoglobins, but they are more likely in individuals where virulent malaria are likely.

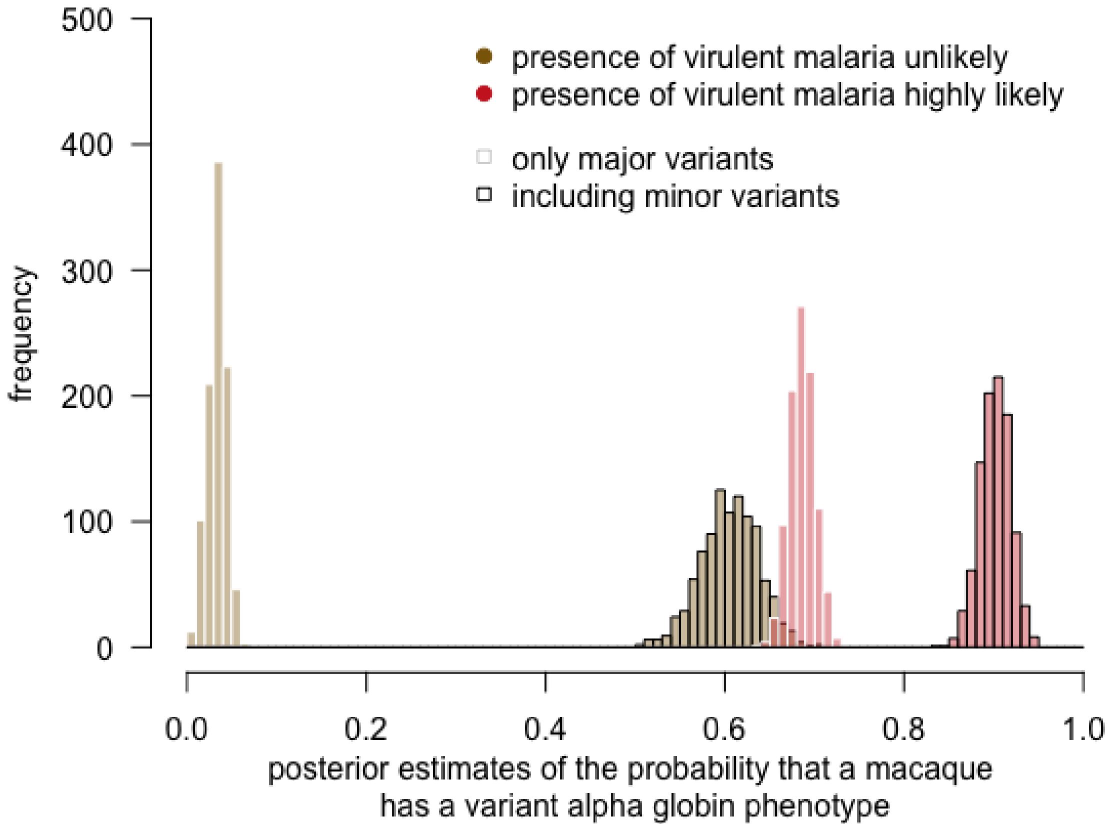

**Supplementary Figure 5. The relationship between the frequency of variant haemoglobins and the heterozygosity of different *M. fascicularis* populations.** Each panel illustrates the relationship between the proportion of *M. fascicularis* exhibiting a variant haemoglobin phenotype (x axis) and observed heterozygosity at the non-globin locus given in the title. For data sources see Supplementary table 4. Haemoglobin phenotypes are as defined in the main text: **A** in the absence of **P** or **Q** is a non-variant alpha globin phenotype, and any phenotype containing **Q**, **P** or both is a variant alpha globin phenotype. Illustrated individual non-globin loci are Albumin (Alb); Acid Phosphatase (AP); NADH dependent diaphorase (Dia); 6 phosphoglucanate dehydrogenase (PGD), phosphohexose isomerase (PHI) and Transferrin (Tf). We also show observed heterozygosity estimated using panels of short tandem repeat (STR) and single nucleotide polymorphism (SNP) loci (Day, et al. 2018). Each marker in each panel is labelled according to location (B=Bali; C=Cambodia; J=Java; M= Malaysia; P=Philippines (unspecified further); P(M)= Philippines (Mindanao); Si=Singapore; Su=Sumatra; T=Thailand). Markers are also classified according to whether or not virulent macaque malarias are present (see legend, Y= yes, N= no, U=unknown). Each panel also displays the Spearman rank correlation coefficient (ρ) and associated p value for each relationship (produced using the ggpubr R package).

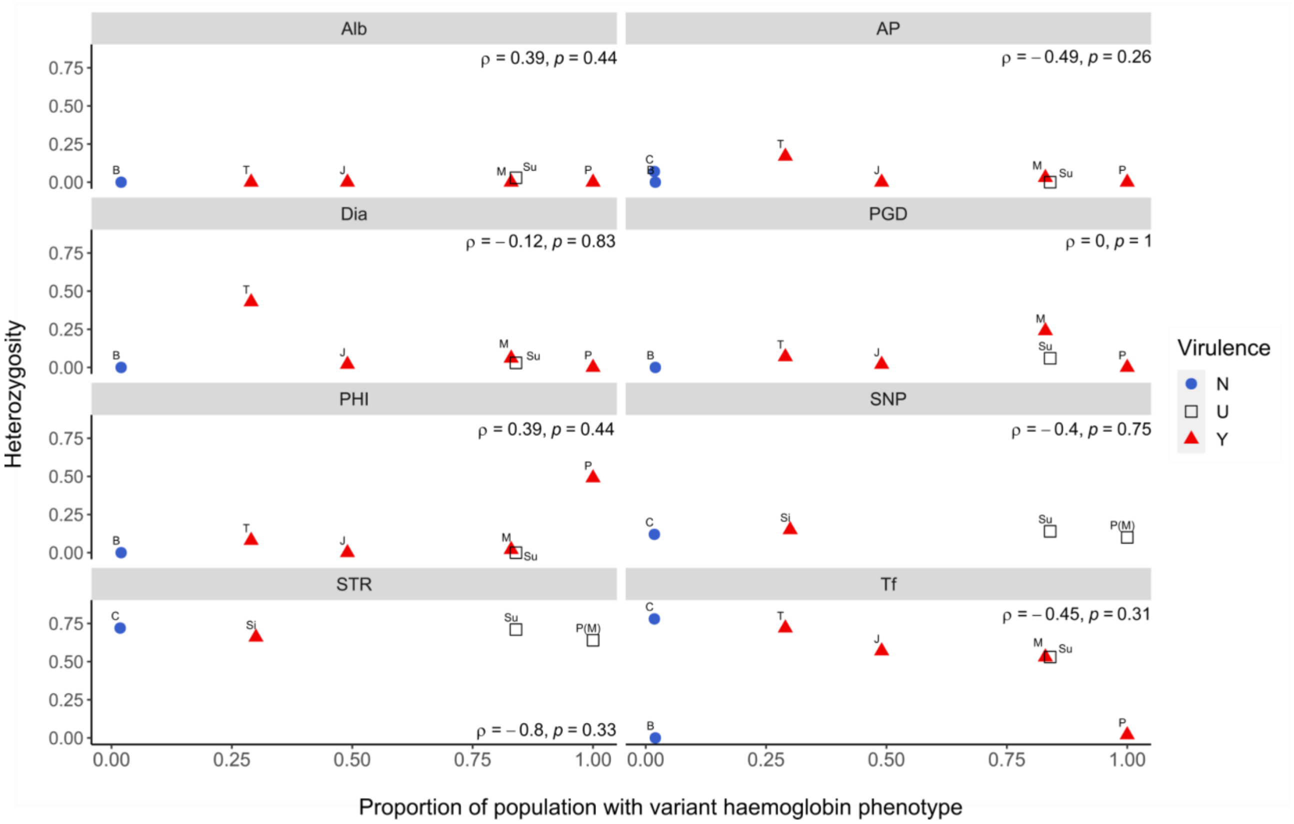

**Supplementary Figure 6. Individual long-tailed macaque *HBA1* sequences.** These stacked bar charts show the frequency of unique sequences found in *HBA1* globin genes of individual macaques. The sequences in red tones (HBA1.3, HBA1.5, HBA 1.10) have a SNP at position 71. Horizontal lines divide the frequencies into 25% intervals (solid) and 16.7% intervals (dashed).

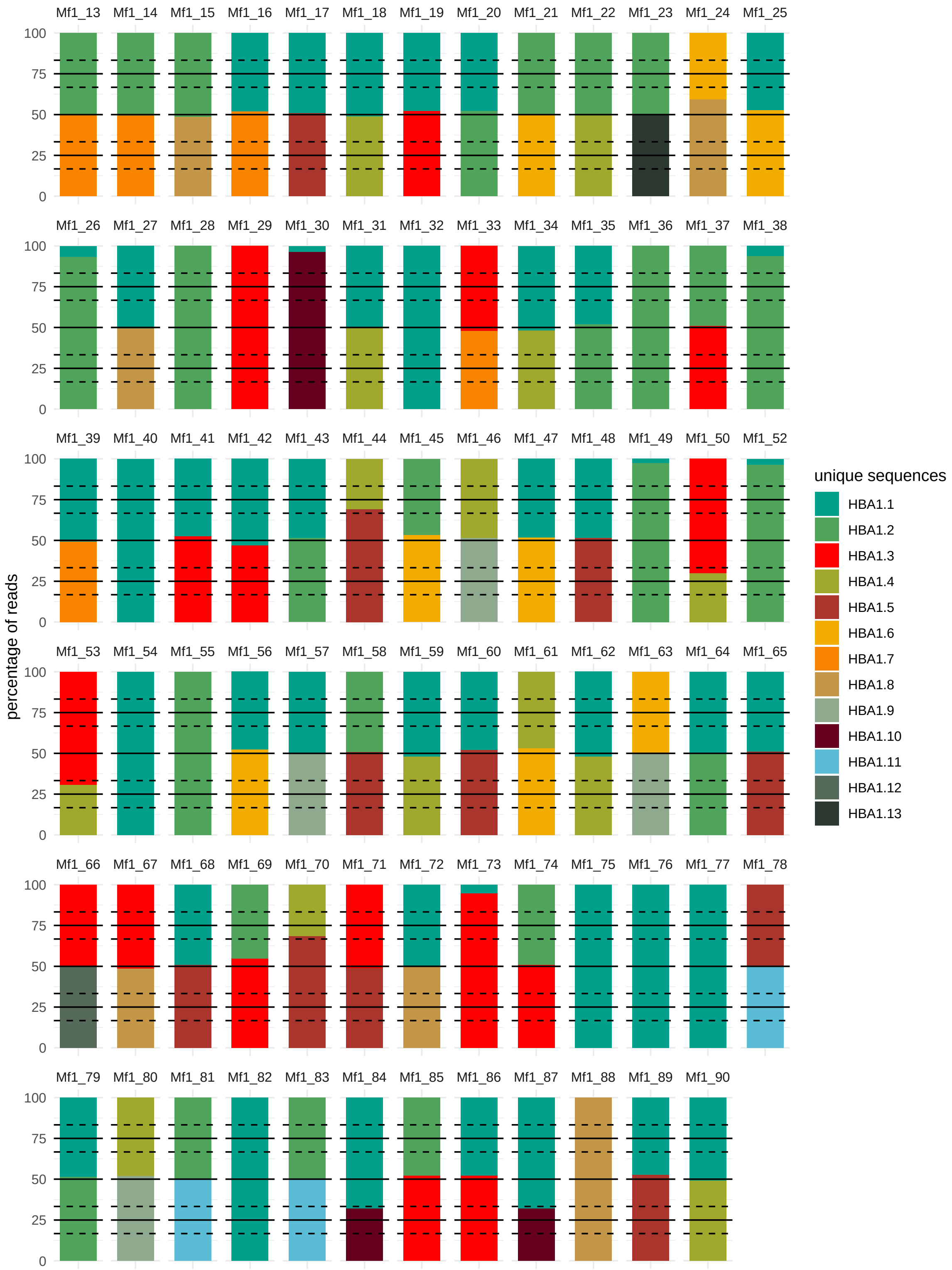

**Supplementary Figure 7. Individual long-tailed macaque *HBA2* sequences.** The stacked bar charts show the frequency of unique sequences found in *HBA2* globin genes of each macaque. The sequences in shades of red (HBA2.1, HBA2.4, HBA2.12) have a SNP at position 71. Horizontal lines divide the frequencies into 25% intervals (solid) and 16.7% intervals (dashed).

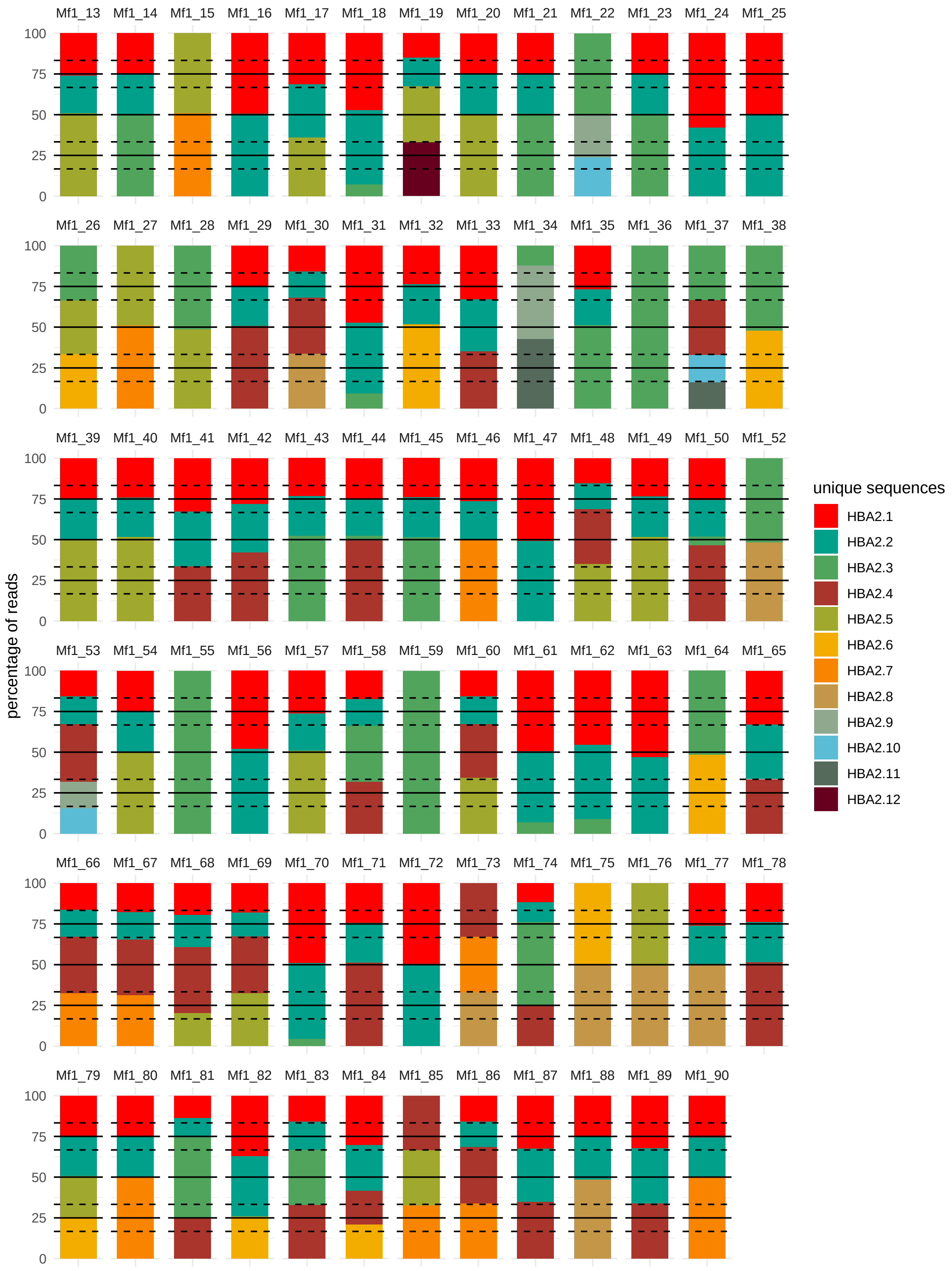

**Supplementary Figure 8. The minimum proportion of any unique *HBA1* or *HBA2* sequence per animal.** Here we visualize the minimum proportions of any unique *HBA1* (x axis) or *HBA2* (y axis) sequence observed in each of the 77 long-tailed macaques. The horizontal and vertical lines indicate the expected minimum proportion of reads possible in an animal with 4, 8, 12 or 20 genomic copies of *HBA1* (vertical lines) or *HBA2* (horizontal lines). Note that 8 copies of *HBA2* per genome could be achieved by 4 copies of *HBA2* on each chromosome, or 2 copies on one chromosome and 6 on another, and so on.

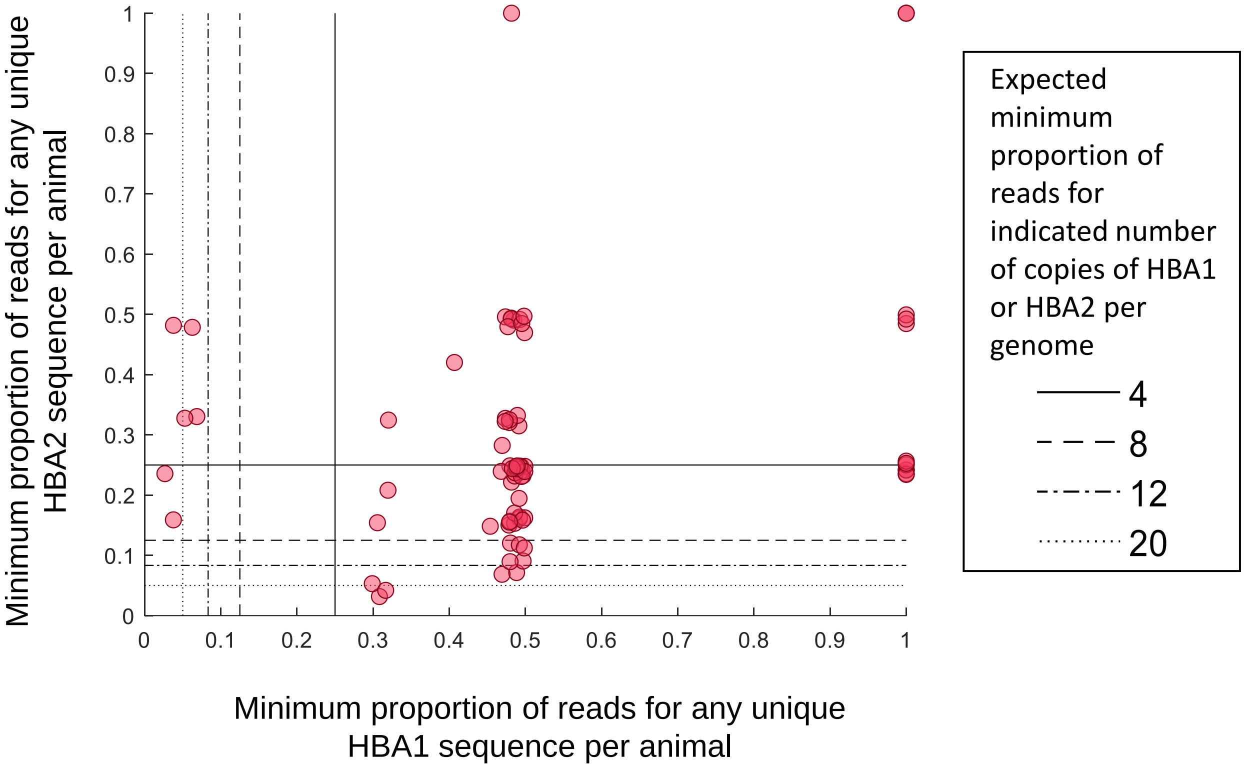

**Supplementary Figure 9. Phylogenetic relationships between *M. fascicularis* alpha globin nucleotide sequences.** A Bayesian phylogeny of alpha globin nucleotide sequences with *P. anubis* as an outgroup. *M. fascicularis HBA1* sequences are indicated in orange text and *M. fascicularis HBA2* sequences are indicated in red text. The regions of *HBA1* and *HBA2* we amplified in this study are found within the *P. anubis* whole genome shotgun sequence (isolate 1X1155) at AHZZ02018146.1:2210-2543 and AHZZ02018145.1:204-537. These sequences are identical, so only one *P. Anubis* sequence is included. We used MrBayes (Ronquist, et al. 2012) to perform the phylogenetic analysis. We used the General Time Reversible model for nucleotide substitution, with gamma distributed substitution rates across different sites, assuming that a proportion of sites are invariable. We fit separate substitution models for non coding regions (the intronic regions up and downstream of exon 2), and for codon positions 1, 2 and 3 within the coding region, meaning that each of these were allowed to evolve at different rates. We ran the MCMC chain for 5,000,000 generations. The consensus tree was visualised using FigTree (Rambaut 2018). Node labels indicate the posterior probability of each split, expressed as a percentage. The scale bar indicates the expected number of substitutions per site. The mean branch length for the split between *P. anubis* and the macaque sequences is 0.0216, with a 95% highest posterior density interval of [0.00788, 0.0385] substitutions per site. Clusters emerge between certain subgroups of *M. fascicularis* sequences, which we have labelled Group A, Group B and Group C. The mean branch lengths for the splits between groups A, B and C and the rest of the sequences are A: 0.003359 [0.000059,0.008361]; B: 0.003462 [0.000006,0.008542]; and C: 0.003236 [0.000006,0.008209] substitutions per site.

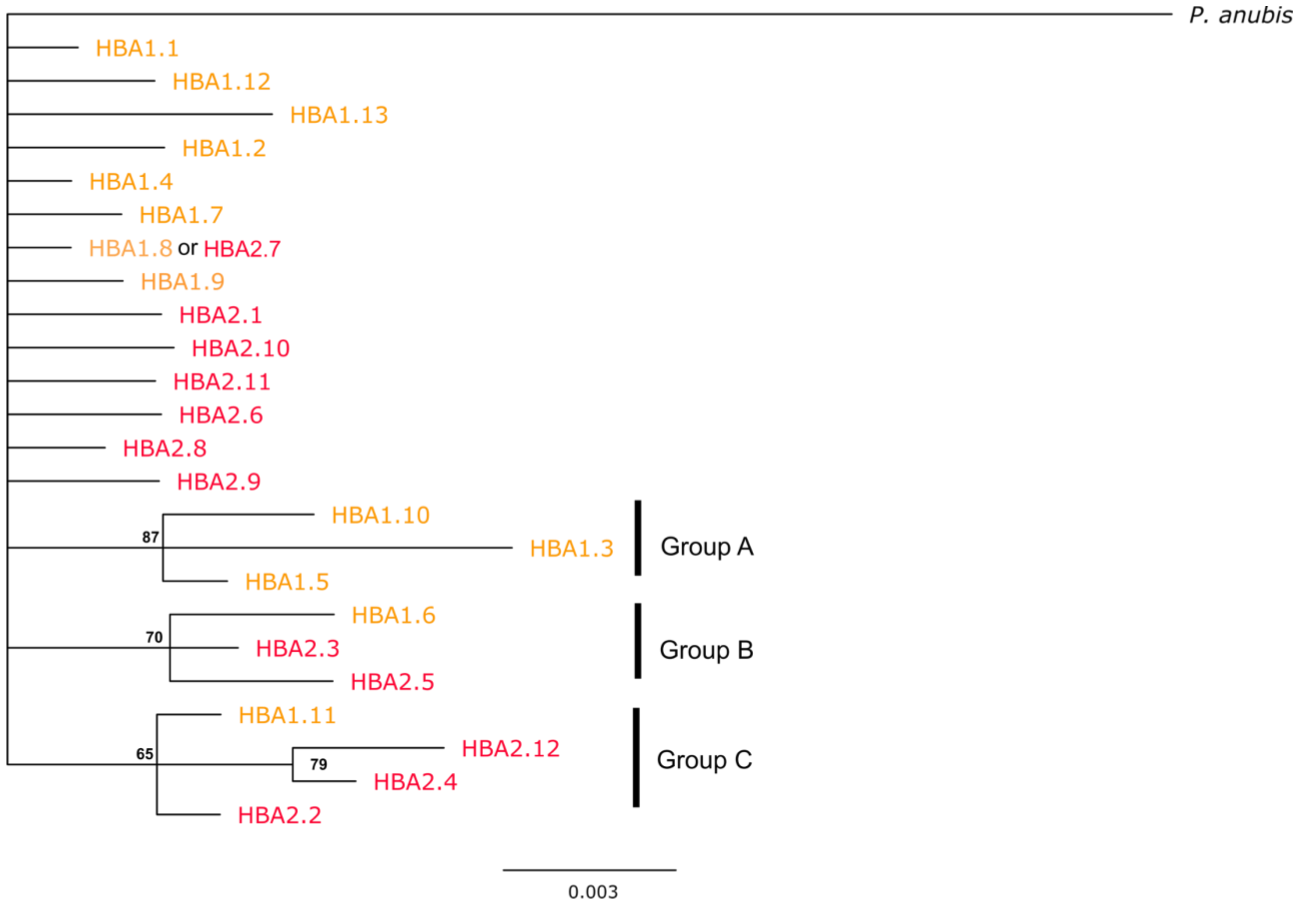

#### Supplemental Information References:


Barnicot N, Wade P, Cohen P. 1970. Evidence for a Second Haemoglobin α-Locus Duplication in *Macaca irus*. Nature 228:379-381.
Barnicot NA, Hewett-Emmett D. 1972. Red cell and serum proteins of *Cercocebus*, *Presbytis*, *Colobus* and certain other species. Folia Primatol 17:442-457.
Barnicot NA, Huehns ER, Jolly CJ. 1966. Biochemical Studies on Haemoglobin Variants of the Irus Macaque. Proc R Soc Lond B Biol Sci 165:224-244.
Barnicot NA, Jolly CJ. 1966. Haemoglobin Polymorphism in the Orang Utan and an Animal with Four Major Haemoglobins. Nature 210:640-642.
Boyer SH. 1972. Extraordinary Incidence of Electrophoretically Silent Genetic Polymorphisms. Nature 239:453-454.
Boyer SH, Crosby EF, Noyes AN, Fuller GF, Leslie SE, Donaldson LJ, Vrablik GR, Schaefer EW, Thurmon TF. 1971. Primate hemoglobins: Some sequences and some proposals concerning the character of evolution and mutation. Biochem Genet 5:405-448.
Buettner-Janusch J, Washington J, Buettner-Janusch V. 1971. Hemoglobins of Lemuriformes. Arch Inst Pasteur Madagascar 40:127.
Choudhury S, Wattal BL, Ramakrishnan SP. 1963. Incrimination of *Anopheles elegans* James (1903) as a natural vector of simian malaria in Nilgris, Madras State, India. Indian J Malariol 17:243-247.
Chuang CH, Lien JC, Lin SY. 1966. Further Studies on Simian Malaria in Taiwan. World Health Organ Bull 567:1-19.
Coatney GR, Collins WE, Warren M, Contacos PG. 1971. The Primate Malarias. Washington DC: US Government Printing Office.
Colless DH. 1956. The Anopheles leucosphyrus group. Trans R Ent Soc Lond 108:37-116.
Day GQ, Ng J, Oldt RF, Houghton PW, Smith DG, Kanthaswamy S. 2018. DNA-based Determination of Ancestry in Cynomolgus Macaques (*Macaca fascicularis*). J Am Assoc Lab Anim Sci 57:432-442.
Dissanaike AS. 1963. *Plasmodium* infection in Ceylon Monkeys. Trans R Soc Trop Med Hyg 57:488-489.
Dissanaike AS. 1965. Simian malaria parasites of Ceylon. Bull World Health Org 32:593.
Dissanaike AS, Nelson P, Garnham PCC. 1965. Two new malaria parasites, *Plasmodium cynomolgi ceylonensis* subsp. nov. and *Plasmodium fragile* sp. nov., from monkeys in Ceylon. Ceylon Med J 14:1-14.
Eyles D, Warren M. 1962. *Plasmodium inui* in Sulawesi. J Parasitol 48:739.
Eyles DE, Laing ABG, Dobrovolny CG. 1962. The malaria parasites of the pig-tailed macaque, *Macaca nemestrina nemestrina* (Linnaeus), in Malaya. Indian J Malariol 16:285-298.
Faust C, Dobson AP. 2015. Primate malarias: diversity, distribution and insights for zoonotic *Plasmodium*. One Health 1:66-75.
Fooden J. 1994. Malaria in macaques. Int J Primatol 15:573-596.
Fungfuang W, Udom C, Tongthainan D, Kadir KA, Singh B. 2020. Stump-Tailed Macaques (*Macaca Arctoides*) are New Natural Hosts for *Plasmodium Knowlesi*, *P. Inui*, *P. Coatneyi* and *P. Fieldi*. Malaria J Under Review.
Garnham PCC. 1966. Malaria parasites and other haemosporidia. Oxford, UK: Blackwell Scientific.
Ho G, Lee C, Abie M, Zainuddin Z, Japnin J, Topani R, Sumita S, Sharm R. 2010. Prevalance of *Plasmodium* in the Long-tailored Macaque (*Macaca fascicularis*) from Selangor, Malaysia. 13th Association of Institutions for Tropical Veterinary Medicine (AITVM) Conference:1-293.
Hoffman HA, Gottlieb AJ, Wisecup WG. 1967. Hemoglobin Polymorphism in Chimpanzees and Gibbons. Science 156:944.
Howard LM, Cabrera BD. 1961. Simian malaria in the Philippines. Science 134:555.
Hrdy DB, Barnicot NA, Alper CA. 1975. Protein polymorphism in the Hanuman langur (*Presbytis entellus*). Folia Primatol 24:173-187.
Huang C-C, Ji D-D, Chiang Y-C, Teng H-J, Liu H-J, Chang C-D, Wu Y-H. 2010. Prevalence and Molecular Characterization of *Plasmodium inui* among Formosan Macaques (*Macaca cyclopis*) in Taiwan. J Parasitol 96:8-15.
Ishimoto G. 1973. Blood Protein Variations in Asian Macaques. III. Characteristic of the macaque blood protein polymorphism. J Anthrop Soc Nippon 81:1-13.
Ishimoto G. 1972. マカク属サルの血液蛋白変異に関する研究. J Anthrop Soc Nippon 80:250-274.
Ishimoto G, Prychodko W. 1970. Hemoglobin Types of Gibbons and Leaf-Monkeys. Primates 11:391-394.
Ishimoto G, Tanaka T, Nigi H, Prychodko W. 1970. Hemoglobin Variation in Macaques. Primates 11:229-241.
Ishimoto G, Toyomasu T, Uemura K. 1968. Intraspecies variations of red cell enzymes and hemoglobin in *Macaca irus*. Primates 9:395-408.
Jeslyn WPS, Huat TC, Vernon L, Irene LMZ. 2011. Molecular Epidemiological Investigation of *Plasmodium knowlesi* in Humans and Macaques in Singapore. Vector Borne Zoonotic Dis 11:131-135.
Kanthaswamy S, Ng J, Satkoski Trask J, George DA, Kou AJ, Hoffman LN, Doherty TB, Houghton P, Smith DG. 2013. The genetic composition of populations of cynomolgus macaques (*Macaca fascicularis*) used in biomedical research. J Med Primatol 42:120-131.
Kawamoto Y, Ischak M, Supriatna J. 1982. Gene constitution of crab-eating macaques (*Macaca fascicularis*) on Lombok and Sumbawa. Kyoto University Overseas Research Report of Studies on Asian Non-human Primates 2:57-64.
Kawamoto Y, Ischak TM. 1981. Genetic Differentiation of the Indonesian Crab-eating Macaque (*Macaca fascicularis*): I. Preliminary Report on Blood Protein Polymorphism. Primates 22:237-252.
Kawamoto Y, Ishida T, Suzuki J, Tanenaka O, Varavudhi P. 1989. A preliminary report on the genetic variations of crab-eating macaques in Thailand. Kyoto University Overseas Research Report of Studies on Asian Non-human Primates 7:94-103.
Kawamoto Y, Tschak TM, Supriatna J. 1984. Genetic Variations Within and Between Troops of the Crab-eating Macaque (*Macaca fascicularis*) on Sumatra, Java, Bali, Lombok and Sumbawa, Indonesia. Primates 25:131-159.

Kitchen H, Eaton J, Stenger VG. 1968. Hemoglobin Types of Adult, Fetal, and Newborn Subhuman Primates *Macaca speciosa*. Arch Biochem Biophys 123:227-234.
Langmead B, Salzberg SL. 2012. Fast gapped-read alignment with Bowtie 2. Nat Methods 9:357.
Lee K-S, Divis PCS, Zakaria SK, Matusop A, Julin RA, Conway DJ, Cox-Singh J, Singh B. 2011. *Plasmodium knowlesi:* Reservoir Hosts and Tracking the Emergence in Humans and Macaques. PLoS Pathog 7:e1002015.
Li H, Durbin R. 2009. Fast and accurate short read alignment with Burrows–Wheeler transform. Bioinform 25:1754-1760.
Matsubayashi K, Sajuthi D. 1981. Microbiological and Clinical Examinations of Cynomolgus Monkeys in Indonesia. In. Kyoto University Overseas Report of Studies on Indonesian Macaque. Kyoto Univ Primate Center Institute. p. 47-56.
Miles A. 2018. pysamstats. Version 1.1.2.
Mulligan HW, Swaminath CS. 1940. Natural infection with *Plasmodium inui* in *Silenus sinicus* from South India. J Malar Inst India 3:603-604.
Nelson P. 1971. *Anopheles elegans,* a natural vector of simian malaria in Ceylon. Trans R Soc Trop Med Hyg 65:695-696.
Nishimoto Y, Arisue N, Kawai S, Escalante AA, Horii T, Tanabe K, Hashimoto T. 2008. Evolution and phylogeny of the heterogeneous cytosolic SSU rRNA genes in the genus *Plasmodium*. Mol Phylogenet Evol 47:45-53.
Nozawa K, Shotake T, Ohkura Y, Tanabe Y. 1977. Genetic Variations Within and Between Species of Asian Macaques. Jpn J Genet 52:15-30.
Otsuru M, Sekikawa H. 1979. Surveys of Simian Malaria in Japan. Zentralbl Bakteriol Mikrobiol Hyg A 244:345-350.
Perwitasari-Farajallah D, Kawamoto Y, Suryobroto B. 1999. Variation in blood proteins and mitochondrial DNA within and between local populations of longtail macaques,on the Island of Java, Indonesia. Primates 40:581-595.
Peyton E, Harrison BA. 1980. *Anopheles* (Cellia) *takasagoensis* Morishita 1946, an additional species in the Balabacensis complex of Southeast Asia (Diptera: Culicidae). Mosquito Systematics 12:335-347.
Prakash S, Chakrabarti SC. 1962. The Isolation and description of *Plasmodium cynomolgi* (Mayer 1907) and *Plasmodium inui* (Halberstadter and Prowazek, 1907) from naturally occurring mixed infections in *Macaca radiata radiata* monkeys of the Nilgiris, Madas State, India. Indian J Malariol 16:303-311.
Putaporntip C, Jongwutiwes S, Thongaree S, Seethamchai S, Grynberg P, Hughes AL. 2010. Ecology of malaria parasites infecting Southeast Asian macaques: evidence from cytochrome b sequences. Mol Eco 19:3466-3476.
Ramakrishnan S, Mohan B. 1962. An enzootic focus of simian malaria in *Macaca radiata radiata* Geoffroy of Nilgiris, Madras State, India. Indian J Malariol 16:87-94.
Rambaut A. 2018. FigTree. Version 1.4.4.
Ronquist F, Teslenko M, Van Der Mark P, Ayres DL, Darling A, Höhna S, Larget B, Liu L, Suchard MA, Huelsenbeck JP. 2012. MrBayes 3.2: efficient Bayesian phylogenetic inference and model choice across a large model space. Syst Biol 61:539-542.
Schmidt L, Greenland R, Rossan R, Genther C. 1961. Natural Occurrence of Malaria in Rhesus Monkeys. Science 133:753.
Seethamchai S, Putaporntip C, Malaivijitnond S, Cui L, Jongwutiwes S. 2008. Malaria and hepatocystis species in wild macaques, southern Thailand. Am J Trop Med Hyg 78:646.
Singh B, Divis P. 2009. Orangutans Not Infected with *Plasmodium vivax* or *P. cynomolgi,* Indonesia. Emerg Infect Dis 15:1657-1658.
Smith D, Ferrell R. 1980. A family study of the hemoglobin polymorphism in *Macaca fascicularis*. J Hum Evol 9:557-563.
Steiper M, Wolfe N, Karesh W, Kilbourn A, Bosi E, Ruvolo M. 2006. The phylogenetic and evolutionary history of a novel alpha-globin-type gene in orangutans (*Pongo pygmaeus*). Infect Genet Evol 6:277-286.
Sullivan B, Nute PE. 1968. Structural and functional properties of polymorphic hemoglobins from orangutans. Genet 58:113.
Takenaka A, Takahashi K, Takenaka O. 1988. Novel Hemoglobin Components and Their Amino Acid Sequences from the Crab-Eating Macaque (*Macaca fascicularis*). J Mol Evol 28:136-144.
Takenaka A, Udono T, Miwa N, Varavudhi P, Takenaka O. 1993. High frequency of triplicated α-globin genes in tropical primates, crab-eating macaques (*Macaca fascicularis*), chimpanzees (*Pan troglodytes*), and orang-utans (*Pongo pygmaeus*). Primates 34:55-60.
Takenaka O, Hotta M, Kawamoto Y, Suryobroto B, Brotoisworo E. 1987. Origin and Evolution of the Sulawesi Macaques 2. Complete Amino Acid Sequences of Seven beta Chains of Three Molecular Types. Primates 28:99-109.
Tomiuk J. 1989. Hemoglobin polymorphism in macaques with reference to the evolution of *Macaca fascicularis* and *Macaca mulatta*. Primates 30:95-102.
Tosi AJ, Coke CS. 2007. Comparative phylogenetics offer new insights into the biogeographic history of *Macaca fascicularis* and the origin of the Mauritian macaques. Mol Phylogenet Evol 42 2:498-504.
Toyomasu T, Ishimoto G. 1969. Transferrin gene distributions in *Macaca irus* from two different local populations. J Anthrop Soc Nippon 77:260-266.
Vythilingam I, Noorazian YM, Huat T, Jiram A, Yusri YM, Azahari AH, NorParina I, NoorRain A, LokmanHakim S. 2008. Plasmodium knowlesi in humans, macaques and mosquitoes in peninsular Malaysia. Parasit Vectors 1:26.
Vythilingam I, Tan CH, Asmad M, Chan ST, Lee KS, Singh B. 2006. Natural transmission of *Plasmodium knowlesi* to humans by *Anopheles latens* in Sarawak, Malaysia. Trans R Soc Trop Med Hyg 100:1087-1088.
Weiss ML, Goodman M, Prychodko W, Moore GW, Tanaka T. 1973. An analysis of macaque systematics using gene frequency data. J Hum Evol 2:213-226.
Xue C, Raveendran M, Harris RA, Fawcett GL, Liu X, White S, Dahdouli M, Deiros DR, Below JE, Salerno W. 2016. The population genomics of rhesus macaques (*Macaca mulatta*) based on whole-genome sequences. Genome Res 26:1651-1662.
Yao L, Witt K, Li H, Rice J, Salinas NR, Martin RD, Huerta-Sánchez E, Malhi RS. 2020. Population genetics of wild *Macaca fascicularis* with low-coverage shotgun sequencing of museum specimens. Am J Phys Anthropol 173:21-33.
Zimmer E, Martin S, Beverley S, Kan Y, Wilson AC. 1980. Rapid duplication and loss of genes coding for the alpha chains of hemoglobin. Proc Natl Acad Sci U S A 77:2158-2162.
